# Spatiotemporally Resolved Multivariate Pattern Analysis for M/EEG

**DOI:** 10.1101/2021.08.17.456594

**Authors:** Cameron Higgins, Diego Vidaurre, Nils Kolling, Yunzhe Liu, Tim Behrens, Mark Woolrich

## Abstract

An emerging goal in neuroscience is tracking what information is represented in brain activity over time as a participant completes some task. Whilst EEG and MEG offer millisecond temporal resolution of how activity patterns emerge and evolve, standard decoding methods present significant barriers to interpretability as they obscure the underlying spatial and temporal activity patterns. We instead propose the use of a generative encoding model framework that simultaneously infers the multivariate spatial patterns of activity and the variable timing at which these patterns emerge on individual trials. An encoding model inversion allows predictions to be made about unseen test data in the same way as in standard decoding methodology. These SpatioTemporally Resolved MVPA (STRM) models can be flexibly applied to a wide variety of experimental paradigms, including classification and regression tasks. We show that these models provide insightful maps of the activity driving predictive accuracy metrics; demonstrate behaviourally meaningful variation in the timing of pattern emergence on individual trials; and achieve predictive accuracies that are either equivalent or surpass those achieved by more widely used methods. This provides a new avenue for investigating the brain’s representational dynamics and could ultimately support more flexible experimental designs in future.

**HIGHLIGHTS:**

- We introduce SpatioTemporally Resolved MVPA (STRM), an approach that explicitly models how successive stages of stimulus processing are distributed in both space and time in M/EEG data.
- We show that STRM is broadly applicable to diverse types of M/EEG data and outputs meaningful and interpretable maps of how neural representations evolve in space and time at millisecond resolution.
- The trial-specific deviations in activity pattern timings identified by STRM are not random, but vary systematically with inter-trial differences in behavioural, cognitive and physiological variables.
- These methods result in predictive accuracy metrics that are mostly equivalent to, or a modest improvement on, conventional methods.

## 1. Introduction

The use of decoding models for the analysis of MEG and EEG data has substantially grown in recent years (Grootswagers et al., 2017). Increasingly researchers are turning to these methods - collectively referred to as Multivariate Pattern Analysis (MVPA) - for their increased sensitivity to distributed patterns of variation that can be attributed to a stimulus (Haynes & Rees, 2006). While projecting high dimensional neural data down to a single metric of classification accuracy affords researchers greater statistical sensitivity, it generally comes at a cost to interpretability: the precise nature of the link between significant changes in decoding accuracy and the underlying neuroscience driving those changes can often be indirect or opaque (Haufe et al., 2014; Kriegeskorte & Douglas, 2019; Naselaris & Kay, 2015; Valentin et al., 2020). Furthermore, commonly used methods offer no sensitivity to patterns that are not perfectly synchronised in time across different trials (Vidaurre et al., 2019).

The main focus in the use of MVPA for M/EEG is to leverage these modalities’ high temporal resolution to investigate the brain’s representational dynamics over time (King & Dehaene, 2014). A convention has emerged for analysing trial data whereby successive spatial filters – each a vector with one linear coefficient applied to each sensor – are trained on data at different timestamps from the presentation of some stimulus. While this convention has certainly been useful, it is not without its flaws, as illustrated in figure 1. Firstly, spatial filters are blind to any temporal structure in the data. These filters cannot detect patterns of temporal autocorrelation that are subtle but consistent over multiple timepoints. Secondly, spatial filters cannot be interpreted as reflecting the brain activity associated with stimuli; this has been widely characterised in the neuroscience literature (Haufe et al., 2014; Kriegeskorte & Douglas, 2019; Valentin et al., 2020), but is somewhat counter intuitive. As shown in figure 1, given an evoked response from two different stimuli in the presence of correlated noise, the coefficients of a linear spatial filter need not visually resemble or reveal these underlying patterns. Methods have been proposed to map backwards from a spatial filter to recover a forward model of the data (Haufe et al., 2014), however these come with a number of caveats – such as a loss of direct interpretability when regularisation is used, as is commonly the case. We instead propose to turn this approach on its head, and ask: why not first learn a generative model of the data, and then use that model to make predictions of unseen stimuli?

**Figure 1:**
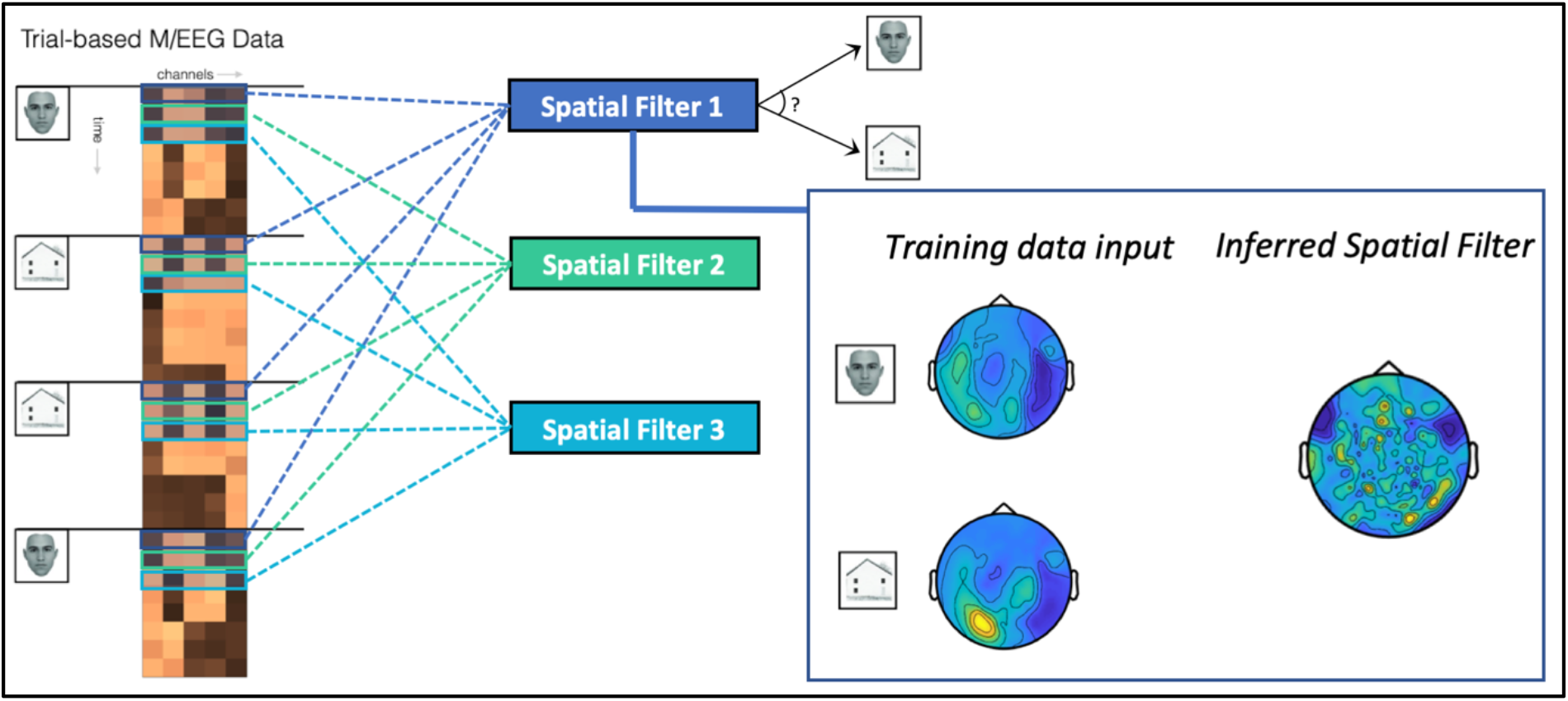
Conventional approaches to decoding in M/EEG are mass univariate through time and difficult to interpret in space. In a typical M/EEG experiment decoding different types of stimuli - in this example, images of faces and houses adapted from (Negrini et al., 2017) (the exemplary maps are based on real MEG data from the dataset presented below) - the conventional approach extracts all data at one timestep from the stimulus onset time and trains a spatial filter on that data to distinguish the conditions. This process is repeated independently at all timesteps. This approach ignores the time series nature of this data, and the spatial filters are neither able to detect patterns that are consistent over multiple timesteps, nor patterns that are consistent over trials but not perfectly aligned in time. It is also challenging for researchers to interrogate why the classifiers made the predictions given; as shown on the right panel and more widely explored in previous literature (Haufe et al., 2014; Kriegeskorte & Douglas, 2019; Valentin et al., 2020), spatial filter coefficients are not directly interpretable with respect to the patterns of activity they are separating, without non-trivial post-hoc analysis.

In neuroscience terminology, this amounts to fitting an *encoding* model, then inverting that encoding model to make predictions through an equivalent *decoding* model (Friston et al., 2008; Haxby et al., 2014; Naselaris et al., 2011). This approach has been successful in fMRI (Casey et al., 2011; Friston et al., 2008; Kay et al., 2008; Mitchell et al., 2008; Naselaris et al., 2009, 2015; Nishimoto et al., 2011; Schoenmakers et al., 2013) but has only seen quite limited adoption for M/EEG (di Liberto et al., 2015; Kupers et al., 2020). In line with these fMRI works, we propose a linear generative model of stimulus evoked activity based upon the popular General Linear Model (GLM) framework. Whilst the GLM in neuroimaging is traditionally associated with mass univariate analysis, it can be extended to be multivariate across space, allowing multivariate predictions to be made on unseen test data through a simple application of Bayes rule (Friston et al., 2008; Haxby et al., 2014). The widespread utility of the GLM approach in neuroimaging suggests it may have a broad applicability across a range of different experimental paradigms; in support of this claim, we demonstrate its use across two very different datasets.

Further, we show how this encoding framework can be extended to take advantage of approaches that adapt to timing differences across different trials (Vidaurre et al., 2019), thereby more fully utilising the high temporal resolution that is the main benefit of M/EEG as a recording paradigm. Extending the work of (Vidaurre et al., 2019), we propose an explicit time series model of the data utilising the Hidden Markov Modelling framework that has seen widespread adoption already in M/EEG analysis (Higgins et al., 2021; Quinn et al., 2018; Vidaurre et al., 2018). In so doing, we are able to characterise the emergence of distinct representational states on individual trials and better model patterns that endure over multiple timepoints but may not be perfectly aligned in time. Previous requirements for patterns to be perfectly aligned over multiple trials limited experimental designs to paradigms that ensure maximal inter-trial reproducibility (Light et al., 2010). Our more flexible modelling approach overcomes this limitation, potentially allowing more ambitious experimental designs in which inter-trial variability – such as that associated with higher cognitive processes – is anticipated and can be quantified. We believe this opens a new door to investigating representational dynamics, by not only characterising when certain aspects of a stimulus emerge in data, but also asking how these representational dynamics are modulated across different trials.

## 2. Methods

Our approach extends the traditional General Linear Model (GLM) framework by (i) incorporating a latent Markov variable to explain time-varying dynamics within trials, and (ii) modelling the multivariate spatial distribution simultaneously over all recorded channels. In so doing, we maintain the benefits of a generative model from the GLM approach, namely interpretable maps of linear activation strengths over each channel; alongside the benefits of multivariate methods, namely an increased sensitivity to distributed patterns of variation over multiple timesteps and amenability to hierarchical modelling (i.e. latent variable modelling of differences in dynamics over trials). These models are fit independently to each subject, resulting in distinct model parameters for each subject upon which standard group statistical tests can be run.

### 2.1. Standard General Linear Model Setup

We begin with the standard formulation of a GLM for evoked response analysis. For a specific subject, with a total of *N* samples of M\EEG data recorded across *P* sensors in a paradigm with *Q* regressors, we denote by *Y* the neural data of dimension [*N* × *P*], with an associated design matrix of regressors *X* of dimension [*N* × *Q*]. This standard GLM setup is not explicit about how the N samples relate to time or trials. The convention in M/EEG analysis is to fit independent GLMs over successive timepoints within a trial in the same manner as in figure 1, such that *N* would equal the number of trials and the effects of each stimulus are modelled independently at each timepoint within the trial. Note that our use of ‘neural data’ here is generic; in this paper we use raw sensor data, but this could equivalently be commonly used features such as bandlimited power. The GLM is formulated as follows:

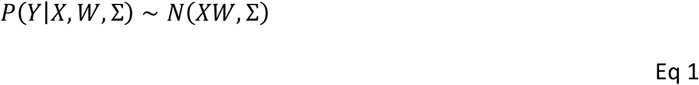

The matrix W, of dimension [*Q* × *P*], is directly interpretable as each row is an estimate of the corresponding stimulus’ activation pattern, while the confidence in that estimate depends on the variance of the corresponding residuals.

The common use of the GLM in mass univariate analysis may lead some readers to mistakenly believe it is a univariate model. This is because, putting aside spatial priors that could be used on *W*, this model stores any multivariate information in the residual covariance matrix Σ; and this matrix is simply not used in mass univariate hypothesis tests that, for example, do group level inference on *W*. However as has been established (Friston et al., 2008), the residual covariance matrix is essential when the model is inverted to make multivariate predictions about unseen test data. Thus, our focus is on the full, multivariate GLM where all terms of the covariance matrix are inferred and used to generate predictions.

### 2.2. SpatioTemporally Resolved MVPA (STRM)

We propose to extend this model hierarchically using the Hidden Markov Model (HMM) framework. This assumes some level of correlation of activation patterns over multiple timepoints that is modelled by a latent state variable, at the same time as having the ability to capture differences in dynamics over trials. Specifically, we model the data vector observed at any point in time *t* on trial *n* as conditional upon a latent, discrete and mutually exclusive state *Z*_*n,t*_ ∈ [1,2, … *K*], as follows:

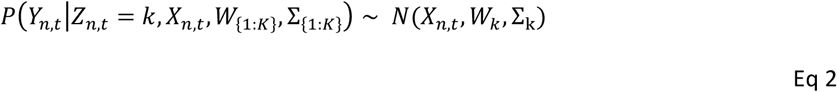

where *K* is the number of states, and the *k*th state is associated with a distinct set of residual covariance patterns Σ_k_ and stimulus activation patterns *W*_*k*_ that linearly map between the trial stimulus X and the data Y. The latent states *Z*_*n,t*_ can then be inferred (with suitable constraints, as outlined below) to explain exactly when each distinct pattern emerges on each different trial, allowing us to infer the combination of activity patterns and corresponding latent state timings that best explain the data at each time point (figure 2).

**Figure 2:**
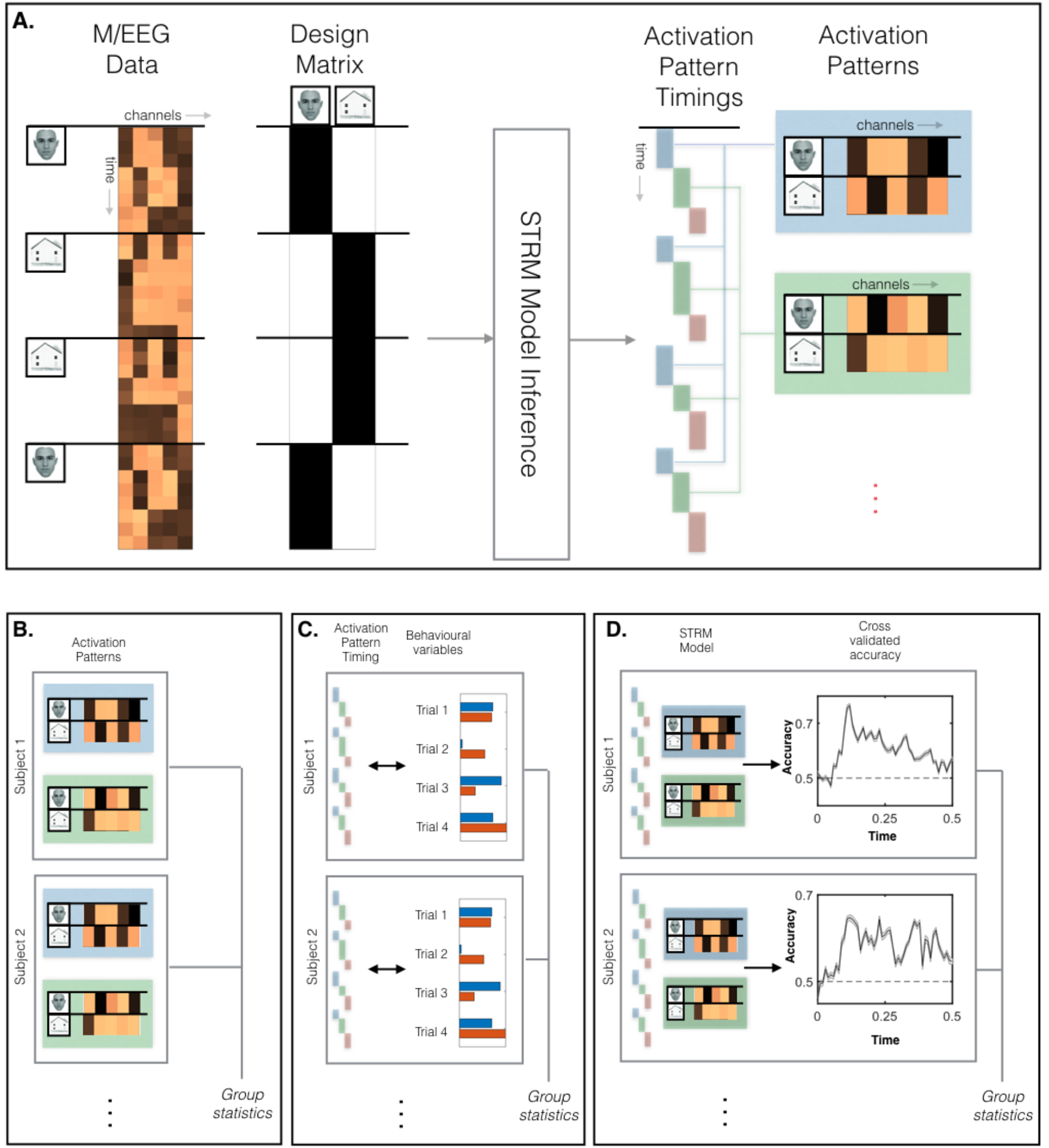
SpatioTemporally Resolved MVPA (STRM) for M/EEG. A. The STRM Model receives as inputs the M/EEG data and corresponding design matrix, and outputs a set of activation patterns and their corresponding timing on each trial. Our analysis pipeline fits this model to data at the subject level, then extracts three different summary statistics (Panel B-D) to analyse at the group level. B. The consistency of state activation patterns can be summarised by taking the group mean activation patterns (as we do for the STRM-Regression model); or of a subject level summary measure such as an ANOVA f-statistic (as we do for the STRM-Classification model). C. Behaviourally meaningful variation in the timing of the activation patterns can be identified by regressing behavioural readouts on individual trials against state timing parameters, and fitting group statistics to the regression parameters. D. The STRM model can be inverted to make multivariate predictions on unseen test data; we can then run standard group statistics on the decoding accuracies obtained.

### 2.3. Bayesian Model Hierarchy

We wish to estimate the unknown model parameters [*Z*_{1:*N*,1:*T*}_, *W*_{1:*K*}_, Σ_{1:*K*}_]. We do this by inferring the full posterior distribution as follows:

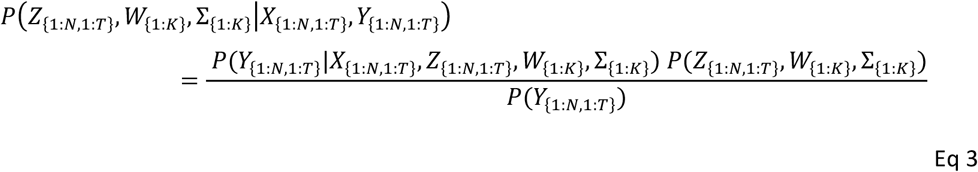

In the below analysis, we expand upon these terms, with modelling decisions explained and justified in turn. We omit the model evidence term (i.e. the denominator in the above equation) as it shall be methodologically sufficient to compute this posterior up to proportion.

#### 2.3.1. Data Likelihood

The likelihood of each observation is conditionally independent of every other observation given the current value of the latent state variable. Thus, we can write the full likelihood of all observations over *N* trials each of *T* timepoints as a product over time and trials:

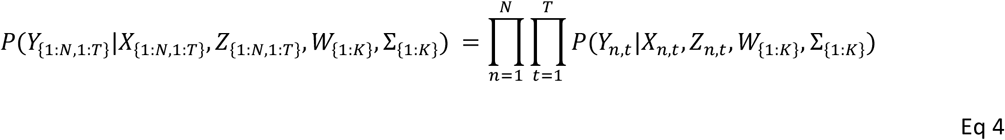

where each individual observation is modelled by a GLM, as in equation 2.

#### 2.3.2. Sequential Latent State Prior

We assume that *P*(*Z*_{1:*N*,1:*T*}_, *W*_{1:*K*}_, Σ_{1:*K*}_) factorises over parameters, i.e.:

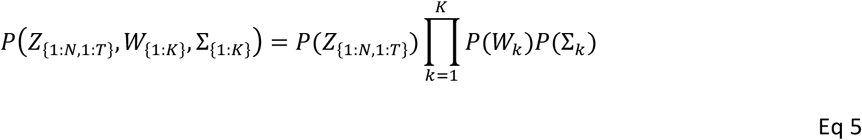

and model the latent state variable *Z*_*n,t*_ using a Markovian prior:

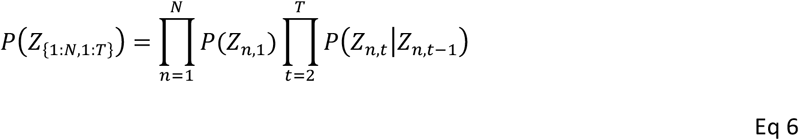

However, we make one important departure from a vanilla HMM. In the cases where HMMs are used for unsupervised analyses of M/EEG data, e.g. to find resting state networks (Higgins et al., 2021; Vidaurre et al., 2018), they are typically allowed to transition freely between states. In supervised data analysis, this can lead to severe overfitting (Ghahramani, 2001), so we instead constrain the model to follow a common trajectory over each trial: we assume that the state order is a fixed sequence, where only the timing of state transitions is allowed to vary. Every trial begins in state 1 and subsequently progresses consecutively through the states, with state transition times freely inferred a trial specific basis (see Figure 3B). This has the advantage of enforcing an interpretable sequence of activation over trials, whilst reducing the number of free parameters governing state transitions. Where unconstrained HMMs must consider a full *K* × *K* transition matrix, this restraint means we need only model a *K* × 1 vector *p*, the *k*th entry of which captures the probability of state *k* transitioning to state *k* + 1. This structure is enforced by the following prior over the state transitions:

**Figure 3:**
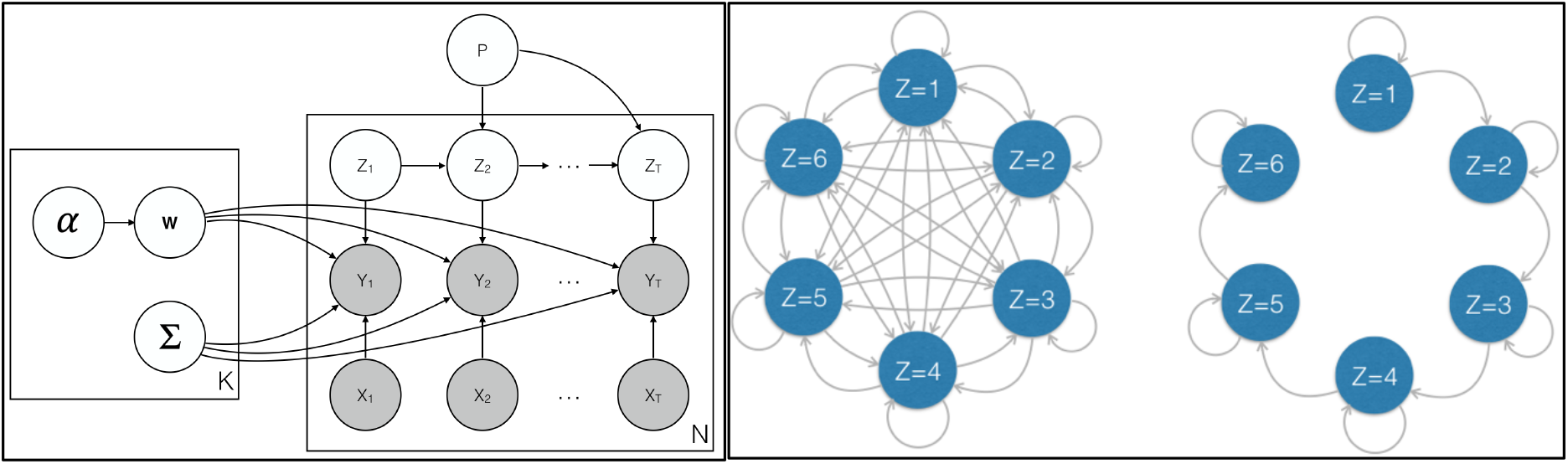
Bayesian model outline and sequential state dynamics. Left Panel: the full model outline in Bayesian plate notation. For each of N trials of length T, we have data observations Y_n,t_ conditioned upon the corresponding design matrix entries X_n,t_. We model these as conditioned upon a latent Markov variable Z_n,t_ which models the state sequence unique to each trial, and upon the associated state parameters W_k_ and Σ_k_, which are modelled separately for each of the K total states. The latent state variables are themselves conditioned upon the transition matrix p, whilst the activation patterns in each state are conditioned upon an automatic relevance determination prior parameter α. Right panel: We depart from conventional HMM modelling, which freely permits any state to transition to any other state as in the diagram on the left, by imposing a sequential latent state prior. As shown on the right, this restricts the permissible state transitions to a consecutive sequence, such that state 1 can either persist or transition to state 2 at each timestep; similarly, if state 2 is active it can either persist or transition to state 3 at each timestep. This structure imposes more aggressive regularisation to overcome the overfitting issues often associated with supervised HMM models.

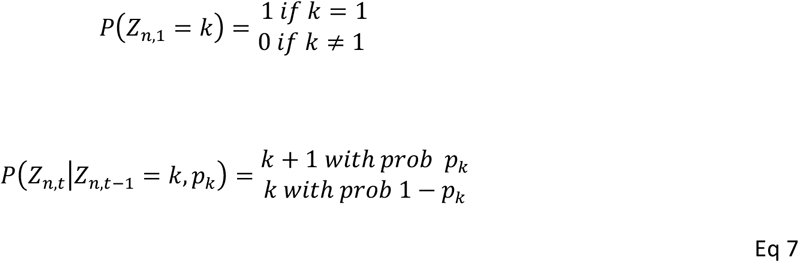

We then set the following conjugate priors over the transition matrix entries:

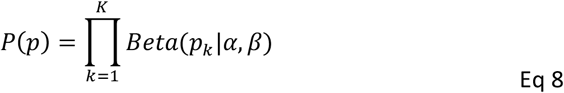

We set the hyperparameters *α* = 1 and *β* = 1 (corresponding to a uniform distribution).

#### 2.3.3. Observation Model Parameter Prior

For the observation model parameters *W*_{1:*K*}_ and Σ_{1:*K*}_, we apply non-informative conjugate priors; this is a Wishart prior for the covariance matrix and an automatic relevance determination (ARD) prior for the stimulus activation patterns. The use of an ARD prior prunes away inferred stimulus activation patterns on sensors that are less consistent over trials, in a manner that performs favourably in neuroimaging (M. W. Woolrich et al., 2009; Yamashita et al., 2008). Denoting by *w*_*k,i,j*_ the *i, j*th entry of the matrix *W*_*k*_, this is implemented as follows:

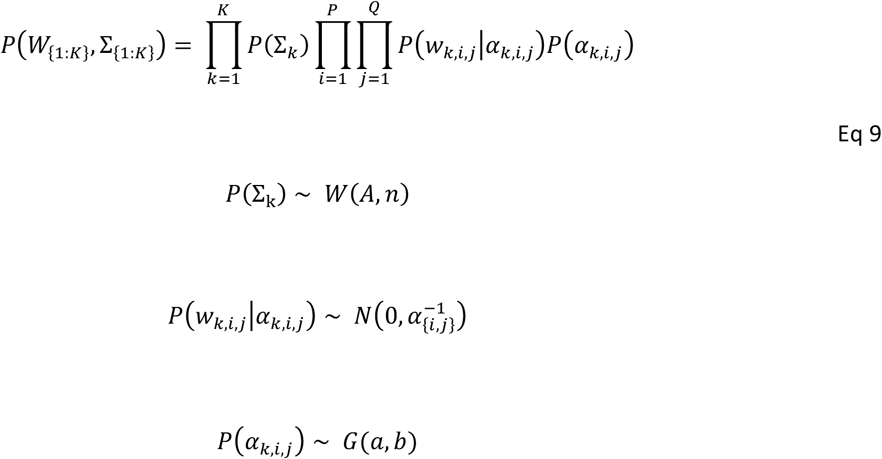

We set the hyperparameters to be non-informative; *A* is set to the identity matrix, *n* = 1, *a* = 1 and *b* = 1. This completes the Bayesian hierarchy.

#### 2.3.4. Variational Inference methods

As a hierarchical extension of a linear model with conjugate priors, this model is amenable to classic variational inference methods. These methods are efficient and scale well to large datasets (Beal, 2003).

### 2.4. Decoding Model Inversion

What we have outlined above is a generative *encoding* model, mapping the spatiotemporal activity patterns associated with each stimulus. Such a model can be inverted to obtain an equivalent *decoding* model by Bayes rule (Friston et al., 2008; Haxby et al., 2014; Naselaris et al., 2011). Specifically, if we define the generative encoding model parameters learned on some training set as *θ* = {*W*_{1:*K*}_, Σ_{1:*K*}_}, for a held-out test set of data *Y* we can make predictions about the associated stimulus *X* as follows:

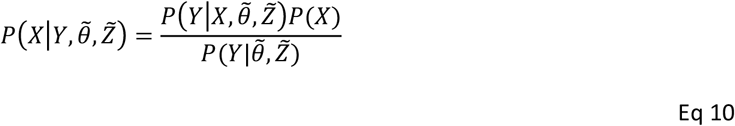

Bayes rule used in this manner allows us to conveniently map between an encoding model and its equivalent decoding model, allowing us to make predictions of the associated stimuli for any unseen test data. Note that 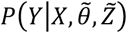 refers to the GLM observation model introduced above (equation 2). When fitting this model to unseen test data, we substitute point estimates 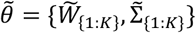 using the expected values from the posterior distributions inferred on the training dataset (equation 9).

We similarly need to estimate the expected values of the latent state variables 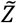 for each timepoint and trial in the test data. Unfortunately, though, as these parameters are specific to each trial, there is no obvious method by which to estimate them for the new, unseen data given *only* the previous training data (Vidaurre et al., 2019). We instead estimate these with a post-hoc procedure (shown schematically in figure S3); taking the training data *Y*_*train*_ and the corresponding posterior state time course probability *P*(*Z*_*train*_), we train a linear regression model at each timestep in a trial to estimate the inferred state timecourses from the data itself: *P*(*Z*_*train*_) = *Y*_*train*_*B*_*t*_ + *ϵ*_*t*_. We then model the test set state timecourse estimate as 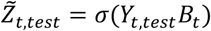, where *σ* denotes the softmax function that ensures 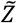 is a mutually exclusive state probability unique to each test set data trial. We discuss the implications of this step later in the paper and expand upon alternative choices in the SI. We also emphasise that this step is only used when computing cross validated accuracy metrics, and as such it does not feature in the initial model fit (figure 2A), the encoding model analysis (figure 2B) or the analysis of pattern timing variation (figure 2C).

The other relevant term for our decoding model inversion is the prior over the structure of the design matrix, *P*(*X*). We consider two cases, the first for where the design matrix consists of mutually exclusive classes (i.e. a classification paradigm); and a second where the design matrix may contain continuous valued regressors (i.e. a regression paradigm). We refer to these as STRM-Classification and STRM-Regression respectively.

#### 2.4.1. STRM-Classification

When the design matrix consists of a total of *Q* mutually exclusive classes, the form of *P*(*X*) is categorical. Assuming the vectors *X*_*t*_ follow a one-hot vector encoding scheme, and writing the prior probability of each class as a [*Q* × 1] vector *C*, we have the following:

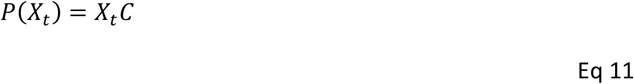

Assuming all classes are equally probable we have all entries of 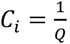.This leads us to an analytical solution to.equation 10 (see Appendix A for details):

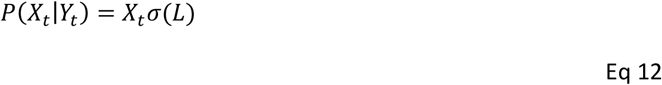

Where *L* is a [*Q* × 1] vector of unnormalized class likelihoods with each entry 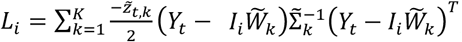; where *I* denotes the [1 × *Q*] vector obtained by taking the *i*th row of the identity matrix of dimension *Q*, and *σ*(*L*) is the softmax function, which outputs a [*Q* × 1] vector with each entry 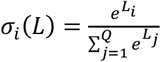. Note that 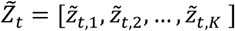 is the [1 × *K*] vector of probabilities of each state being activated at time *t*; these are the terms discussed above that must be estimated by a regression model.

This solution is equivalent to classification by Linear Discriminant Analysis (LDA); so, with the inferred latent state dynamics, 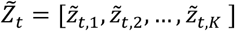, we now have a dynamic form of LDA. Note that in different applications, users may also choose to model the covariance matrix Σ as strictly diagonal; in which case this model is equivalent to a dynamic form of Gaussian Naive Bayes classification.

#### 2.4.2. STRM-Regression

Similarly, given a model that relates a set of continuous valued regressors *X*_*t*_ to observed data *Y*_*t*_, we can make new predictions given some assumption of the overall distribution of the regressors; we will assume they are standardised and uncorrelated, such that:

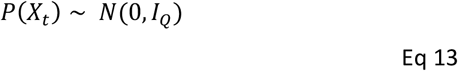

where *I*_*Q*_ is the identity matrix of dimension *Q*. As a conjugate prior, this ensures for any new observation 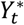 that the posterior of equation 10 has a Gaussian distribution (see appendix B):

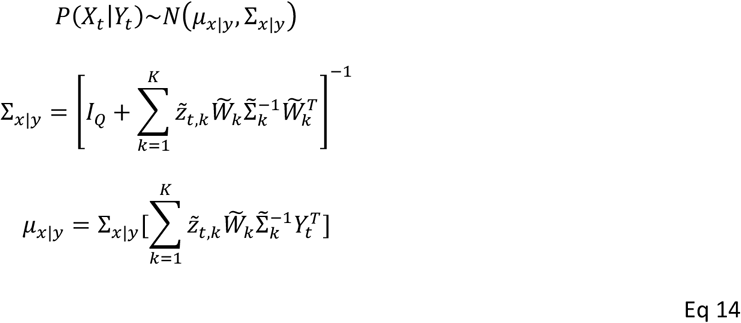

In the absence of the latent state variable, 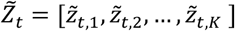, this is a case of Linear Gaussian Systems (LGS) model (Murphy, 2012; Roweis & Ghahramani, 1999); thus, with the inclusion of the latent state variable, we have a dynamic LGS model.

### 2.5. MEG Visual Data Analysis

The first dataset we analyse comprises a visual stimulus decoding task previously published as part of a larger study (Liu et al., 2019).

#### 2.5.1. Task outline

All participants signed written consent in advance; ethical approval for the experiment was obtained from the Research Ethics Committee at University College London under ethics number 9929/002. A total of 22 participants fixated on a cross onscreen and were presented with visual stimuli in a randomised order. There was a total of eight distinct visual stimuli (for details see (Liu et al., 2019)). To ensure continuous engagement with the task, on 20% of trials the stimuli were inverted; the participant was required to push a button to indicate if the stimuli was inverted. The below analysis uses only the data on the non-inverted trials.

#### 2.5.2. Data preprocessing

MEG data was acquired at a rate of 600 samples per second on a 275 channel CTF scanner. Data was filtered within a passband of 0.1 - 49Hz, downsampled to 100 samples per second using a polyphase low-pass filter with cutoff 25Hz, then epoched to extract periods in time from the moment of stimulus presentation to 500 milliseconds later. To ensure training data was balanced across classes, for each subject we determined the number of trials in each class and, if *n* is the number of trials of the least sampled class, only kept the first *n* trials of all other classes. This resulted in balanced class sets for each subject, with different subjects having between 26 and 33 total samples of each class. At this point the data was separated into two streams. First, the sensor space data to be used for learning the decoding models underwent dimensionality reduction using PCA with eigenvalue normalisation, keeping the top 50 components (which accounted for 98.2% of the total variance) (Grootswagers et al., 2017). Second, and separately, the same data was projected into source space using a LCMV beamformer (Veen et al., 1997; M. Woolrich et al., 2011) projecting onto an 8mm MNI grid. Source data was then parcellated into 38 anatomically defined Regions of Interest (ROIs) derived from an independent component analysis of fMRI resting state data from the Human Connectome Project (Colclough et al., 2016). Source leakage was then corrected for by orthogonalization as outlined in (Colclough et al., 2015).

#### 2.5.3. STRM-Classification Model

As the stimulus set was categorical, we established a design matrix with 9 regressors; these corresponded to one regressor for each distinct visual stimulus, along with an intercept term. Thus, the inferred model coefficients for each latent state corresponded to a generic activation pattern over all stimuli for that state (i.e., the intercept term), and an effect specific to each visual stimulus for that state. We initially fixed the parameter for the number of states to *K* = 8 and sought to explore the activation parameters and state timing information for this fixed number of states. Only later in the pipeline, when determining classification accuracy, do we seek to optimise this parameter.

We fit our STRM-Classification model to the principal component space derived from the sensor level data as in (Grootswagers et al., 2017). Models were fit independently to the data from each subject. When evaluating spatial activity maps and state timecourse variability, we fit a single model to all data for each subject; when evaluating classification accuracy we trained and tested an ensemble of models for each subject in a cross-validation procedure outlined below.

#### 2.5.4. Characterising spatial activity maps

Having determined the state timings from fitting our STRM model, we projected the state timing information back onto the original sensor data to obtain sensor activity maps per stimulus per state, and then for additional interpretability onto the equivalent source localised data to obtain source space activity maps per stimulus, per state and per subject (Figure 2B). As we are interested in the regions that *differentiate* the different stimuli, we computed the f statistic for a within subject ANOVA at each ROI, effectively asking how different the representations of the eight stimuli were at that particular ROI, for that subject. Maps in figure 4 plot the average f-statistic over subjects per voxel, thresholded at the top 75th percentile for visualisation.

**Figure 4:**
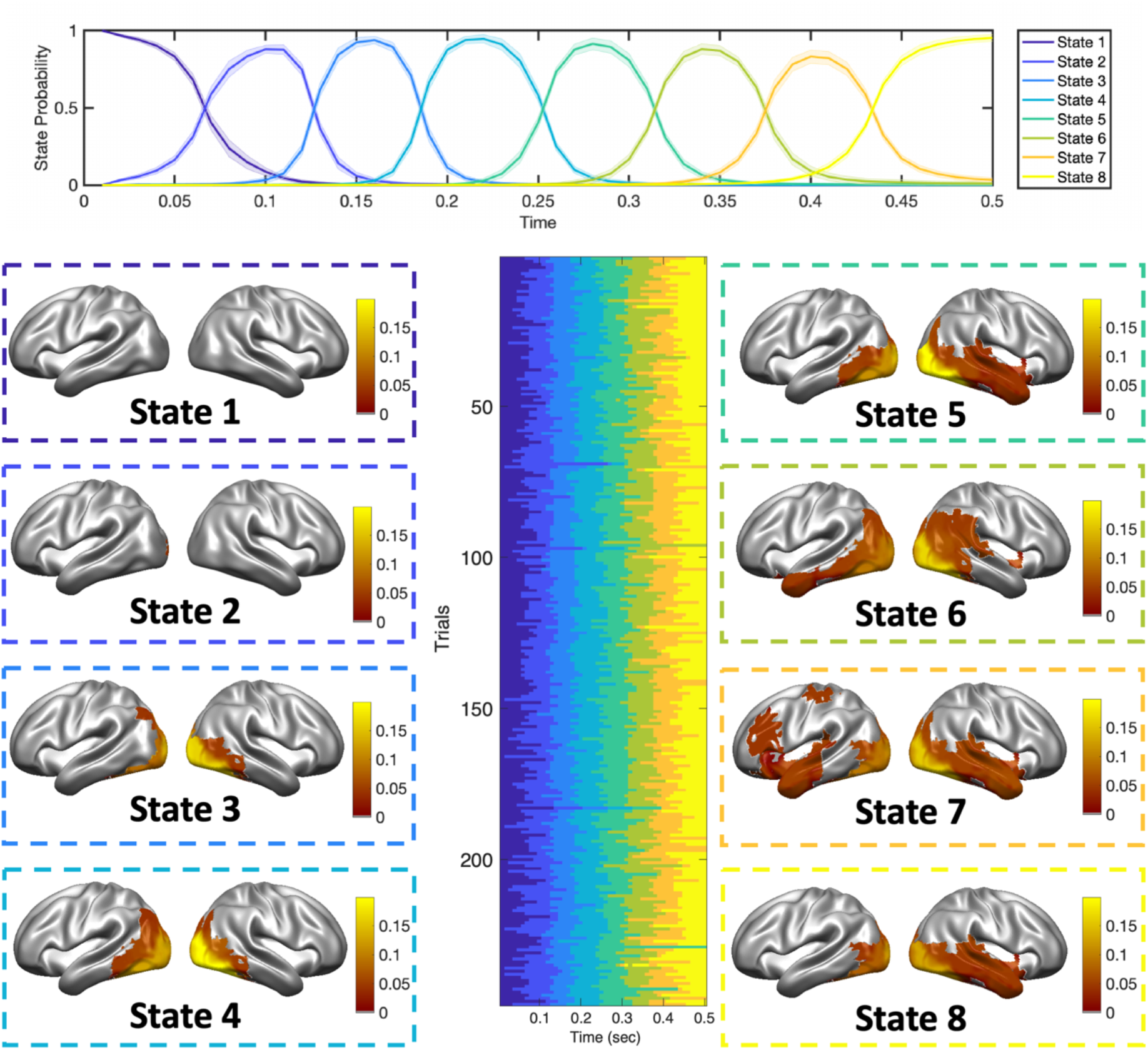
Resolving the successive stages of visual stimulus processing in space and time. Fitting the STRM-Classification model independently to each subject’s MEG data recorded during visual stimulus presentation, using K = 8 states allows us to investigate the stages of visual stimulus processing and the times on individual trials at which they emerge. Top panel: Average timing of each state over a trial (mean +/- ste over subjects), demonstrating the mean time after stimulus presentation that each state emerges. Lower central panel: a raster plot of state timings inferred for a sample subject over 248 trials, demonstrating the variability in timings over successive trials within the common sequential pattern progressing from state 1 to 8. Lower outer panels: the thresholded, group-mean f statistics, per ROI, as a result of multiple subject-level ANOVAs; this displays the amount of information contributed by that ROI to discriminate the different visual stimuli (see Methods). Statistics are thresholded at the 75th percentile of all test statistics obtained.

#### 2.5.5. Characterising state timecourse variability

The state time courses inferred from fitting our STRM model characterise the timing at which specific activity patterns emerge on individual trials. We wished to explore what variables might influence these timings, and so we fit a post-hoc multiple regression model asking whether specific behavioural and physiological variables allowed us to predict the timing of specific states on individual trials (as illustrated schematically in Figure 2C).

For each subject, given a STRM fit with *K* states, we fit (*K* − 1) multiple regression models, with the *k*th model predicting the timing of the transition from the *k*th to the (*k* + 1)th state (i.e. using the state transition time as the dependent variable). In each model we used the same four independent variables; the first two regressors were the inter-stimulus interval (ISI) and participant reaction time (see figure 5), capturing variation in the overall timed structure of the trial. Note that the inter stimulus interval was randomly generated for each trial from a uniform distribution between 0.5 and 2 seconds, whereas the reaction time was determined by the participant and thus reflects a behavioural readout.

**Figure 5:**
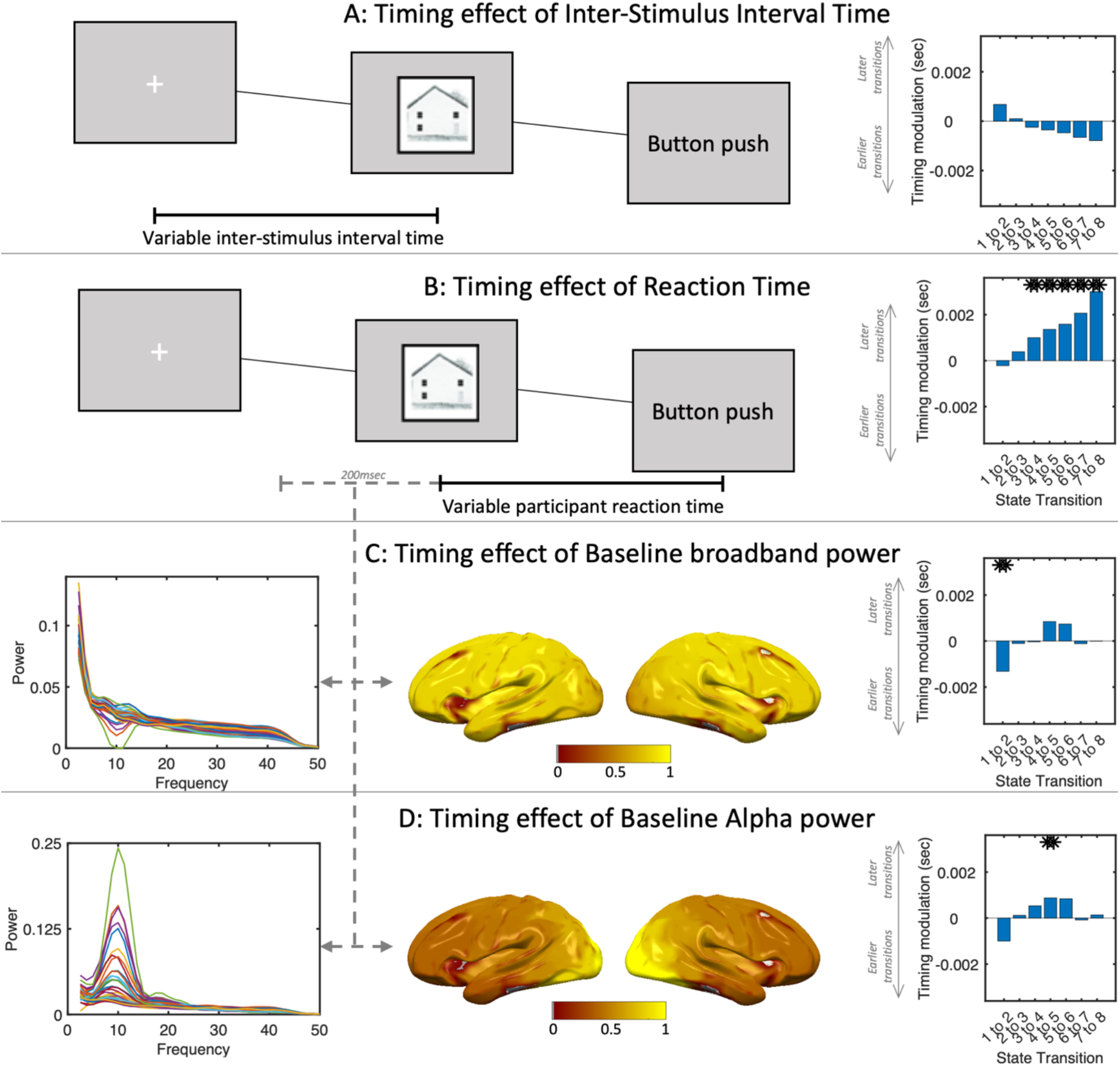
The timing of visual processing is modulated by behaviour and physiology. The timing of visual processing is modulated by behaviour and physiology. Panel A: inter-stimulus intervals do not significantly modulate state transition times. Panel B: Longer participant reaction times are predictive of delayed transitions into states 3 to 8, with increasing effect size towards later states. Panel C: increases in baseline broadband power are associated with more rapid transitions into state 2, an early visual processing state. Panel D: increases in baseline alpha power over visual areas is associated with delayed transitions from state 4 into state 5. In all bar plots, asterisks denote significance at Bonferroni corrected levels (p=2.1e-3).

The remaining two independent regressors were designed to determine whether spontaneous changes in baseline electrophysiological patterns immediately prior to image presentation influenced the speed at which different patterns emerged. For this, we computed the power spectral density (PSD) in each ROI of the source space data over the 200msec preceding stimulus presentation. This data was considerably high dimensional, so we applied a non-negative matrix factorisation across the data from all subjects to extract 2 primary modes of spatio-spectral variation. Specifically, we arranged the baseline PSD estimates for each subject across $N$ ROIs in *F* frequency bands and *M* trials into a single matrix of dimension [*M* × *NF*]. We then applied a Non-Negative Matrix Factorisation (NNMF) decomposition to the row-wise concatenation of all subjects’ PSD matrices as in (Vidaurre et al., 2018), obtaining 2 main modes of spatio-spectral variation that corresponded to a broadband mode and a visual alpha mode (see figure 5). We used the expression strengths of each of these two modes on each trial as the two independent variables in the multiple regression model, thus asking the degree to which baseline broadband power and baseline visual alpha power influenced the timing of subsequent visual processing states.

Finally, to eliminate collinearity we decorrelated all four regressors using symmetric orthogonalization. These multiple regression models were fit at the subject level and comprised a total of 4(*K* − 1) multiple comparisons. We evaluated significance of effects at the group level through two tailed t-tests on the distribution of model coefficients over subjects, using Bonferroni correction of p values.

#### 2.5.6. STRM-Classification Predictive Accuracy

To assess the performance of STRM-Classification model, we used ten-fold cross validation. This entailed a ten-fold partitioning of each subject’s data, followed by an iterative procedure of holding out one fold for testing while training the model on the remaining data for that subject, then testing the classification performance using the procedure outlined in equation 10 on the held-out partition. This was repeated iteratively until all trials had been classified; we defined classification accuracy as the proportion of trials upon which the correct label was predicted by the classifier.

Whereas we previously held the parameter *K* for the number of states fixed to a single value, the classification accuracy provides a clear metric by which this parameter could be optimised. Thus, the above cross validation procedure was run using a total of ten different values of *K* ranging from 4 to 22 in steps of 2. These can either be evaluated separately or optimised by subject level cross validation. This latter procedure entails holding out one subject, determining the value of the parameter *K* that maximises the classification accuracy for the remaining subjects, and selecting that value when determining the accuracy for the held-out subject.

To assess statistical significance, we used two tests. To test whether the STRM-Classifier accuracy was significantly greater than the equivalent timepoint-by-timepoint classifier accuracy as a function of time, we used non-parametric cluster permutation tests with a t threshold of 1 (Nichols & Holmes, 2003). To test whether the overall accuracy (averaged over all timepoints) exceeded other classifiers, we first applied a group level ANOVA followed by pairwise t-tests.

### 2.6. EEG Data Analysis

To demonstrate the general applicability of this method, we also applied it to a set of EEG data collected during a decision making and reward learning task. This dataset was selected as it used stimuli (i.e. reward outcomes) that varied continuously rather than categorically, corresponding to the STRM-Regression model. Furthermore, the more complex task design involved additional variables that we believed may modulate neural processes and thus state timing patterns.

#### 2.6.1. Task outline

A total of 30 participants completed the task whilst undergoing EEG scanning. 23 were of sufficient data quality to be included in the analysis (the other participants were excluded either because of not understanding the task or excessive noise). The task consisted of navigating between different foraging patches in an effort to maximise the total reward accumulated over the course of the recording session. Specifically, there were two types of decisions participants had to make. First, for the ‘site selection’ phase, they chose between one of two patches; they then entered their chosen patch (after a variable waiting period), and accrued reward (the ‘Reward accrual’ phase). During the reward accrual phase, participants were shown their current reward level which changed every second (i.e. decaying from different starting points and different decay rates). Participants were continually prompted: they could either do nothing and continue to accrue reward at a depleting rate, or press the space bar to collect their accumulated reward during this patch visit. They would then again be asked to choose between the two patches. Decay rates and starting points of both patches were learnable but changed periodically throughout the experiment.

For the purposes of our analysis, the only task event analysed is the period where participants receive the overall reward (the ‘Reward receipt’ phase). At that moment, participants had to estimate the average reward rate they had achieved in the current trial in order to decide to leave earlier or later in the next trial or indeed pick the other patch next time. The average reward rate is computed by combining the accumulated reward at a patch and the time invested to receive it.

All participants signed written consent in advance; ethical approval for the experiment was obtained from the Medical Sciences Inter-Divisional Research Ethics Committee at the University of Oxford under project number R28535/RE001.

#### 2.6.2. Data preprocessing

Electroencephalographic signals were recorded with active electrodes from 59 scalp electrodes mounted equidistantly on the standard 10-20 system elastic map (EasyCap). All electrodes were referenced to the right mastoid electrode and re-referenced offline to the average. Continuous EEG was recorded using a SynAmps RT 64-channel Amplifier (1000 Hz sampling rate). The data were epoched from -1 second to 2 second around the reward events. The data were then band-pass-filtered at [1-35] Hz. We denoised our EEG data in a multiple step procedure. First, we removed trials containing large artefacts semi-manually artefact using ’ft rejectvisual’ function in FieldTrip. This function visualises trial variances and allows the user to remove trials and electrodes directly from the GUI. We did this so that the following ICA would not be dominated by very few excessive noise trials. We then ran an ICA and correlated the component timecourses with the computed vertical and horizontal EOG’s. If an ICA component showed high correlations with either EOG and had a characteristic topography for eye movements or blinks, we regressed out that component. In the next step we removed trials with excessive Muscle artefacts by taking eight electrodes (’PO7’, ’PO8’, ’POZ’, ’PO4’, ’PO3’, ’O1’,’OZ’, ’O2’) near the neck which were most affected by muscle movements, z-scoring them and removing trials with a conservative z-score cutoff at 20. After muscle filtering a second visual inspection was applied to the data in case some very noisy trials remained. We downsampled the data to 100 samples per second using a polyphase filter with 25Hz cutoff. Additionally, to further remove ocular artefacts, we regressed the EOG channel signals from each EEG electrode signal. We applied dimensionality reduction as previously described for the MEG data, using PCA with eigenvalue normalisation and keeping the top 20 components (out of 59 channels), which explained a total of 93.6% of the variance.

#### 2.6.3. STRM-Regression Model

Each trial consisted of a presentation on screen of the reward value that had just been accumulated. For our STRM-Regression model, we used a design matrix, *X*, with 2 continuous regressors: the first being a mean activation value to model the mean change in activity for a state over all trials, and the second being the value of reward presented on screen in that trial. This second regressor was normalised over trials.

We fit the STRM-Regression model to the principal component space derived from the sensor level data as in (Grootswagers et al., 2017). This is done independently to the data from each session for each subject (note that for any group statistics we first averaged model parameters for a given subject over both of their recording sessions, then ran group statistics on these subject average values). As outlined in section 2.5.3 we use a total of *K* = 8 states (as previously, we initially hold this parameter fixed while characterising the model output, and later show how it can be optimised for decode accuracy).

#### 2.6.4. Characterising spatial activity maps

Having determined the state timings from initially fitting STRM to the principal component space derived from the sensor level data, we then projected the state timing information back onto the original sensor data to obtain sensor activity maps per regressor per state. Maps in figure 7 plot the group average value of model coefficients for each state; that is, the mean activity pattern associated with each state and the value evoked effect within each state.

#### 2.6.5. Characterising state timecourse variability

In exactly the same way as conducted for the MEG data, having determined the state timings from fitting our STRM-Regression model, we then asked whether the timings of these stimulus processing states were significantly modulated by other variables within the task - specifically, variables that the participant is required to track in order to optimally complete the task.

For each subject, we fit (*K* − 1) multiple regression models, with the *k*th model predicting the timing of the transition from the *k*th to the (*k* + 1)th state (i.e., using the state transition time as the dependent variable). In each model we used three independent variables. These consisted of the value of the stimulus itself; the total time invested at the site (i.e. how long the participant had spent in order to accumulate the value shown in the stimulus); and the recent reward history (i.e. what the participant might expect to gain from leaving the site in search of another). To eliminate collinearity these variables were decorrelated by symmetric orthogonalization and normalised (Colclough et al., 2015). After fitting these models to the data for each subject, we conducted a group analysis testing whether any of the regressor coefficients were significantly different from zero. This involved 3(*K* − 1) multiple comparisons; p values were evaluated for significance after Bonferroni correction.

#### 2.6.6. STRM-Regression Predictive Accuracy

In the same manner as outlined for the MEG data, we then assessed the performance of the STRM-Regression model to predict continuous stimuli using ten-fold cross validation and showed how the parameter *K* controlling the number of states could be optimised by cross-validation over subjects. We used as accuracy metric the Pearson correlation between predictions and the true values. To assess statistical significance, when comparing accuracy vs time of two classifiers we applied a cluster permutation test with t-stat threshold of 1 (Nichols & Holmes, 2003); when comparing the overall accuracy (averaged over all timepoints) of a group of models we applied a group ANOVA.

## 3. Results

We present here the results obtained from fitting the same model to two separate datasets. The first is a set of MEG recordings of categorical visual stimulus presentations, the second a set of EEG recordings of participants completing a cognitive task.

### 3.1. Decoding Visual Stream Representations using MEG

Visual stimuli evoke a cascade of feedforward and feedback activity through the dorsal and ventral visual streams (Goodale & Milner, 1992; Hochstein & Ahissar, 2002). Existing methods to identify the spatiotemporal evolution of these representational dynamics have relied upon methods that fuse MEG and fMRI recordings (Cichy et al., 2014) or require invasive recording types (Goodale, 1993). We asked whether these results could be corroborated using MEG as the sole recording modality, by utilising our STRM-Classification model. We fit our STRM-Classification model to a previously published dataset comprising MEG recordings of 22 participants viewing randomly presented visual images from a set of 8 stimuli (Liu et al., 2019).

#### 3.1.1. STRM Visual Stream Activation Patterns

Given STRM-Classification models fit independently to each subject’s data, we asked how consistent the inferred spatial patterns of activity were over subjects. Specifically, we asked whether ROIs emerged that were particularly informative of class differences, and whether these were the same across subjects. Nothing in the model expressly enforces common patterns of activation for a given state across subjects, these are only unified by their common position in the temporal sequence of states (see figure 3). Figure 4 plots the mean state sequence timings across all subjects for a STRM-Classification model fit with K=8 states. As demonstrated in the central panel of the figure, which shows the inferred state timecourses on individual trials for an example subject, all trials follow the enforced sequential state sequence whilst allowing for variability in precise activation pattern timing on individual trials.

The STRM-Classification model was fit to the principal component space computed from the sensor data (see methods); to assess which ROIs were most informative as to the different class, we projected the state timecourses onto the source space data and conducted subject-level ANOVAs to determine which ROIs displayed the greatest variation over conditions. Plotting the group-mean f statistic for each ROI identifies a clear progression through the visual hierarchy. Applying a percentage threshold across all states (see methods), state 1 identifies no significant sources of information, consistent with its interpretation as preceding the arrival of visual information to cortex. State 2, which accounts for periods around 100msec after stimulus onset that show elevated decoding accuracy, similarly shows no single ROI above thresholding, suggesting that these early visual representations may have limited spatial consistency over subjects. States 3 and 4 show the propagation of this information from occipital visual cortex to outer visual areas; these states encompass periods in time 120 to 250msec after stimulus onset. Throughout all subsequent states, the visual cortex remains the dominant source of information relevant to the decoding, however from states 5 to 8 significant information additionally emerges in inferior temporal and then lateral temporal areas. Thus, information first appears in temporal areas in state 5 which corresponds to roughly 250msec after stimulus presentation, then endures to the end of the recording 500msec after stimulus presentation.

#### 3.1.2. Activation Pattern timings are modulated by behaviour and physiology

Our STRM model resolves the specific timings on each individual trial when patterns of stimulus-associated activity emerge. We then asked whether these timings were meaningful – specifically, how they related to other measures that varied over individual trials. We investigated this by fitting a multiple regression model to predict the timings of the transitions between different states using the state timecourses inferred by STRM. This multiple regression (fit independently for the transition times between each pair of states) had two regressors indicating event timing on each trial (reaction times and ISIs), and two regressors indicating spontaneous variations in broadband power and visual alpha power during a baseline period 200msec prior to stimulus presentation (Sederberg et al., 2019). This tested whether behaviour and/or changes in the baseline distribution of power affected the timing of visual information processing.

We found a strong relationship between response times and state transition times. This is significant for all states from 3 onwards, with an increasing effect size towards later states. ISIs, on the other hand, did not exhibit a significant relationship; there is an apparent trend whereby longer ISIs were associated with slower state sequences, but this was not significant after multiple comparison correction.

Spontaneous changes in the baseline power similarly modulate the subsequent propagation of visual processing states in a manner that appears selective to specific stages in the visual hierarchy. High levels of broadband power are associated with earlier transitions out of state 1 into state 2, whereas high levels of visual alpha band power are associated selectively with later transitions out of state 4 into state 5 – the state we associate with emergence of information outside visual cortex. This is broadly consistent with proposed roles for alpha governing top-down control of attention and gating of information as it moves downstream (Jensen & Mazaheri, 2010); the relationship between broadband activity and earlier visual states is perhaps more surprising, however such patterns have been shown to be strong modulators of neural activity in a wide variety of ways (Miller et al., 2014; Podvalny et al., 2019).

These examples serve primarily as a model validation, demonstrating that the observed variation in activity pattern timings does not arise randomly but rather is reflective of underlying neural processes that are behaviourally and physiologically relevant. They furthermore suggest that activation timings in different stages of the visual cortex are not modulated uniformly, but rather that different variables may selectively affect propagation timings in different parts of the visual hierarchy.

#### 3.1.3. Classification Accuracy

An advantage of MVPA is that model fit can be straightforwardly assessed using the metric of classification accuracy. We therefore asked how STRM-Classification compared to conventional analyses. Figure 6A plots the classification accuracy obtained by STRM-Classification versus that of the equivalent conventional approach, namely timepoint-by-timepoint LDA classification. This model does lead to enhanced classification accuracy, over later timepoints (where activity patterns might be expected to be less well aligned); however, this gain is relatively modest. It appears that although the state timings correlate with behavioural and physiological information, they do not result in dramatic gains to classifier performance. Performance is reasonably robust to the STRM model order; as shown in the second plot of Figure 6, performance is equivalent for *K* = 10 or higher. This plot furthermore identifies a performance tuning curve, making this amenable to parameter optimisation and motivating the cross-validation approach we use to generate the upper and lower panels of Figure 6 (see methods).

**Figure 6:**
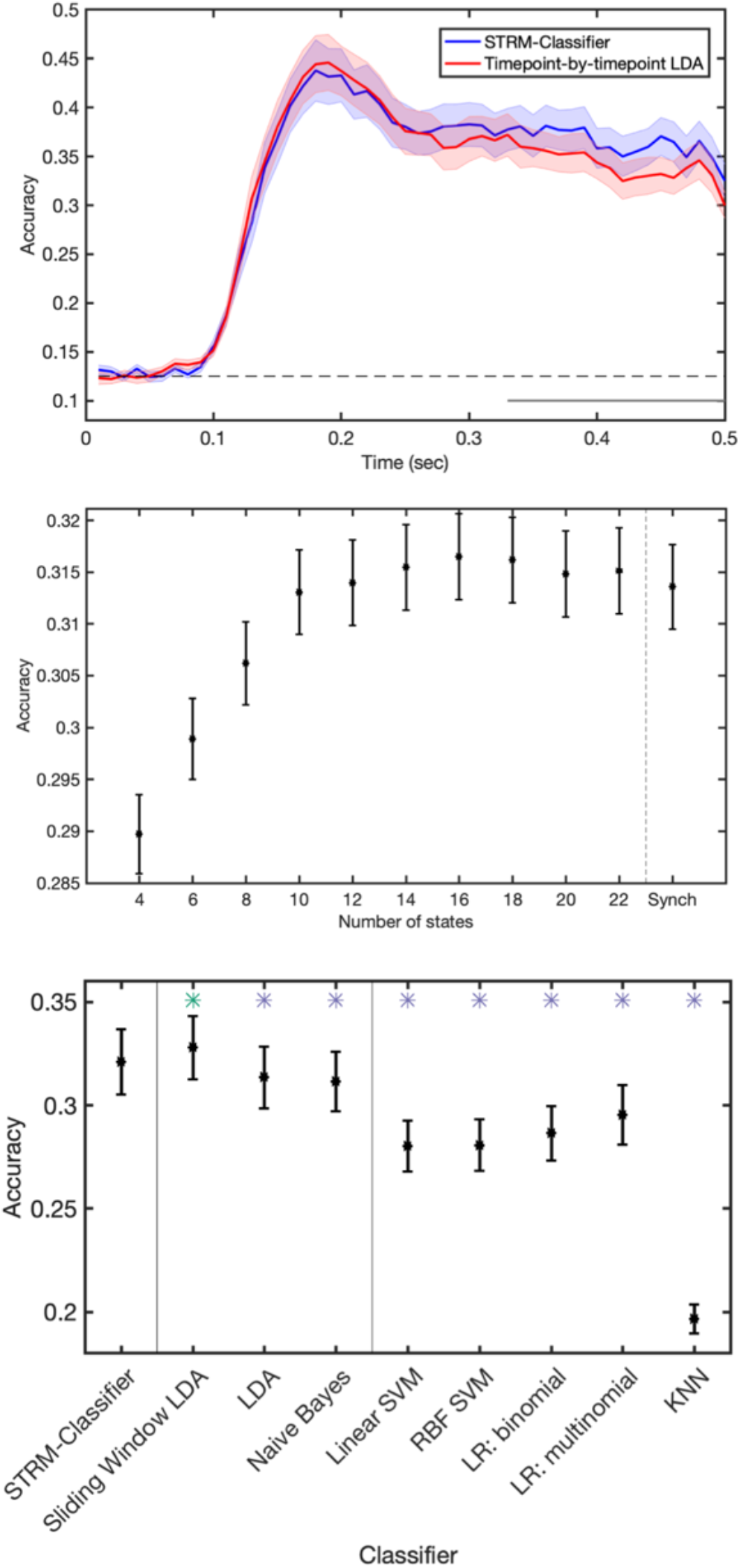
Classification accuracy achieved by different MVPA methods. Top panel: Plotting the accuracy (mean over subjects +/- ste) vs time for STRM-Classifier (with K optimised through cross validation – see methods) vs timepoint-by-timepoint spatially resolved decoding identify marginal improvements in classification accuracy over later timepoints when temporal patterns are identified. Middle panel: Plots of mean accuracy over all timepoints between 0-0.5 seconds as a function of the number of states K; plot shows mean over subjects +/- ste. This shows this relationship is robust for values of the parameter K above a sufficient level; equivalent decoding accuracy is achieved when using K = 10 states or higher. Lower panel: the STRM-Classifier performs favourably when compared to discriminative classification methods. We here compare the STRM classifier with 8 other classification methods – three other spatially resolved classifiers (LDA with optimised sliding window length - see methods; LDA using the conventional timepoint-by-timepoint decoding approach, and Naïve Bayes using the conventional timepoint-by-timepoint decoding approach); and additionally 5 different discriminative classifiers fit using timepoint-by-timepoint methods (Linear SVMs, non-linear SVMs using a radial basis function (RBF) kernel; binomial and multinomial logistic regression (LR), and K-nearest neighbour (KNN) classifiers with K optimised through cross validation). We find in general that generative encoding model based classifiers (STRM, LDA and Naïve Bayes) outperform discriminative classifiers (SVM, LR and KNN), and furthermore that STRM decoding outperforms equivalent methods when used with the conventional timepoint-by-timepoint decoding approach; however, we similarly find that these gains are slightly surpassed by optimised sliding window methods. Asterisks denote significantly different from STRM-Classifier accuracy at Bonferroni corrected levels; green asterisks show significantly higher accuracy (Sliding Window LDA); blue significantly lower (all other classifiers).

**Figure 7:**
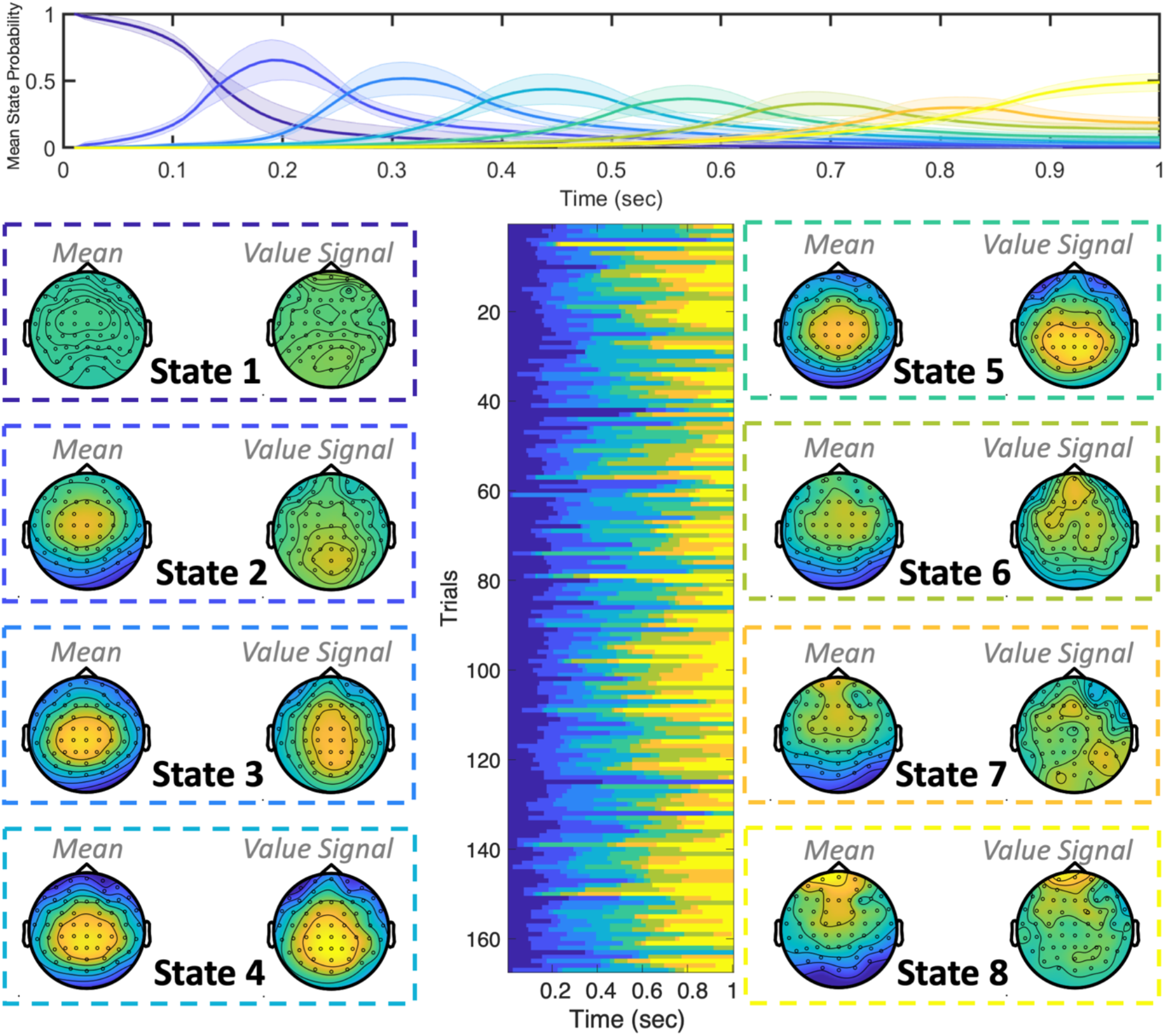
The stages of EEG Value Processing. Fitting the STRM-Regression model independently to each subject’s EEG data with K=8 states identified a consistent pattern of sequential activity on each trial, but with significant variation in the timing of events on individual trials. Top panel: mean state timecourse +/- ste over subjects. Lower centre panel: example state timecourses for one subject exhibiting significant variation over trials. Outer panels: Mean (over subjects) activation patterns for each state and each regressor. Each trial is associated with consistent patterns of activity, comprising a mean pattern of activation common to that stage of cognitive processing on all trials and a separate value-specific component. Both components are characterised by medial parietal activation; in the case of the value signal this appears to emerge initially in parietal areas (for example, state 2) and later move to more frontal regions (state 6). In response to concerns about eye movement artefacts accounting for significant decode-ability, none of these topographies appear eye movement related.

There are two innovations that this performance could be ascribed to; it could be due to the forward modelling procedure and the assumptions it imposes (which in this case are equivalent to the assumptions of LDA); and/or it could be due to the time-varying dynamics that our model permits. To test the effect of the forward modelling procedure, we compared the accuracies achieved by a range of different classification methods. First, we compared the classification accuracy obtained using generative classifiers (LDA and Naïve Bayes) with those achieved by a range of discriminative machine learning classifiers (Support Vector Machines – linear and with radial basis function kernels; logistic regression in binomial and multinomial configurations; and K-nearest neighbour classifiers). All classifiers were implemented on a timepoint-by-timepoint basis. As shown in Figure 6, generative model-based classifiers offer significant improvements in classification accuracy over these methods. Thus, we conclude that – at least for this dataset – the assumptions imposed by our generative model accurately represent the data and afford greater classification accuracy as a result.

### 3.2. Decoding Stages of Cognitive Processing in EEG

The ability to discern and analyse the timing of activity patterns emerging across different trials is potentially more salient in the case of non-sensory representations. Recently, decoding has emerged as a popular paradigm for analysing higher cognitive functions in complex tasks at high temporal resolution (Holdgraf et al., 2017; Kriegeskorte & Kievit, 2013), however these methods still mostly assume these higher cognitive functions are perfectly aligned across trials. We thus asked whether the model we had proposed generalised to a very different EEG dataset, that involved prediction of a continuous variable (value) over multiple trials in a complex foraging task, where contextual variables differed substantially across trials and could potentially determine activity pattern timings.

#### 3.2.1. Spatially resolved stimulus activation maps

Each trial within the foraging task consisted of participants being presented with an amount of reward they had just accumulated (see Methods and Figure 8 for full task outline). We first set out to decode the amount of reward presented on each trial and determine the origins of the activation patterns that predict it.

**Figure 8:**
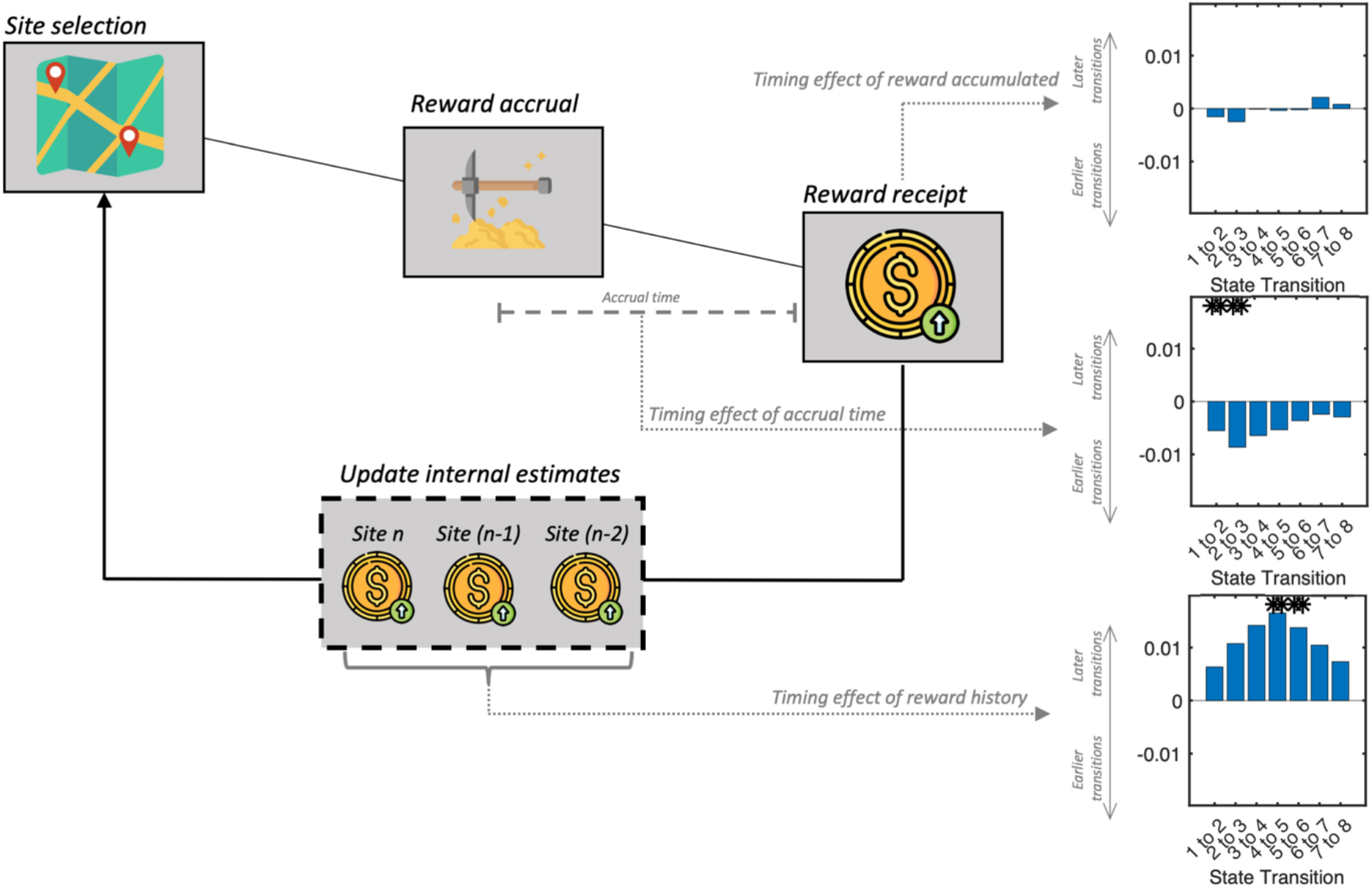
The timing of value processing is modulated by cognitive variables. We investigated whether latent cognitive variables within the overall task structure significantly affected the timing at which different stages of value processing emerged on different trials. Specifically, we asked whether three regressors – the overall reward accumulated, the accrual time, and the recent reward rate – could predict the transition times between states on individual trials. We found no significant effect of reward accumulated, the variable being decoded, however we found significant effects of accrual times (reflecting how much time the participant invested to obtain the reward) and recent reward history (reflecting the expected value of alternative sites). Longer accrual times were associated with more rapid transitions through early stimulus processing states, whereas high rewards in recent history were associated with delayed transitions into intermediate stimulus processing states.

We fit STRM-Regression models, independently for each subject and session, to one-second epochs of EEG data following presentation of the amount of reward the participant had received. This allowed us to identify clear sensor space spatial topographies associated with each regressor in the STRM-Regression design matrix, with the first regressor representing a common mean pattern for each state over all trials and a second representing the value of the reward received. As shown in Figure 7, these maps identify patterns emerging initially over medial parietal regions in sensor space and propagating forward over successive states. The spatial location of the value signal is consistent with the literature on value encoding, which is associated with encoding initially in the superior parietal cortex and later in the medial prefrontal cortex (Hunt et al., 2012; Kolling et al., 2012). Whilst the sensor space maps in Figure 7 do not afford the same precision in terms of anatomical interpretation as achieved in the visual MEG source space maps of Figure 4 (note that we did not perform source reconstruction on the EEG data due to the moderate variability in cap and electrode positioning across subjects), they nonetheless suggest an anatomically derived interpretation: we can loosely interpret states 1-4 as representing the stages of value processing concentrated on parietal areas, whereas states 5-8 represent the emergence of a value processing state more concentrated on medial frontal regions.

One common concern using decoding in EEG is whether the prominent artefacts associated with eye movements may confound the decoding accuracy. To make neuroscientific statements, experimentalists must be assured that the underlying source of their decoders’ performance is neural and not some muscle or ocular confound. Common decoding techniques are unable to indicate the role that such artefacts may have played in the predictions made by the decoder; whilst it is common to provide a post-hoc analysis of ERPs to refute such claims, this evidence can only ever be indirect as it is does not explain how the classifiers made the predictions. In contrast, the predictions made by this method are a measure of how closely the spatial patterns match the parameters shown in figure 7; the clear midline activity patterns that these maps show are therefore direct evidence to refute claims of muscle confounds and eye motion artefacts driving the decoding result.

#### 3.2.2. Timing of value processing is modulated by cognitive variables

Effective completion of the foraging task requires participants to reconcile the amount of reward they receive on each trial with (i) the ‘accrual time’, i.e. the amount of time they had invested at the site in order to accrue that much reward, and (ii) what they would currently expect to receive if they moved to a different patch. For example, a given stimulus has a different intrinsic value if the participant had spent a particularly long time to receive it; or if the participant expected to accrue reward more quickly elsewhere. These two variables we will refer to as cognitive variables, as they are not explicitly presented to the participant on screen as a stimulus but must be estimated and tracked alongside the completion of the task.

We asked whether these cognitive variables influence the timing at which value is processed on individual trials. We investigated this by fitting a multiple regression model to predict the timings of the transitions between different states using the state timecourses inferred by STRM. This multiple regression had three regressors: accumulated reward, accrual time and recent reward rate. We found no significant relationship between value (the regressor being decoded) and state transition times at Bonferroni corrected levels. We did find a strong relationship between accrual time and state progression, with longer times invested at the site associated with more rapid transitions into states 2 and 3, the very earliest stages of stimulus processing in parietal cortical areas. We similarly found a relationship with the recent reward history: on trials where much more reward had been received recently, the onset of intermediate stages of stimulus processing (states 5 and 6 specifically) were delayed. Successful completion of this task requires participants to discount the value of a stimulus against how much time they invested to receive it and how it compares to recent trials; these results suggest these two comparisons are undertaken at different times and specifically modulate the timing of distinct stimulus processing states.

Once again, these results serve fundamentally as model validation; they demonstrate that the inferred timing of value processing on individual trials varies systematically with cognitive variables that reflect key parameters within the task. Notably, again this timing is modulated differentially by the different regressors – that is, early stages of stimulus processing are influenced by accrual times whereas intermediate stages of stimulus processing are modulated by the recent reward rate. Viewed alongside the anatomical maps of each state, these differential effects could suggest distinct variables modulating the parietal and prefrontal components of value processing.

#### 3.2.3. Accuracy of Dynamic LGS Decoding

Finally, we asked whether the enhanced sensitivity afforded by STRM-Regression resulted in benefits to decoding accuracy. We measured accuracy by the cross validated Pearson Correlation between test set model predictions and their true values. We found no significant differences in performance between STRM-Regression and other metrics. We also asked how sensitive this behaviour was to the parameter controlling the number of states; as shown in figure 9B it is very consistent across all values of this parameter tested (no significant differences were observed; one way ANOVA p=0.98).

**Figure 9:**
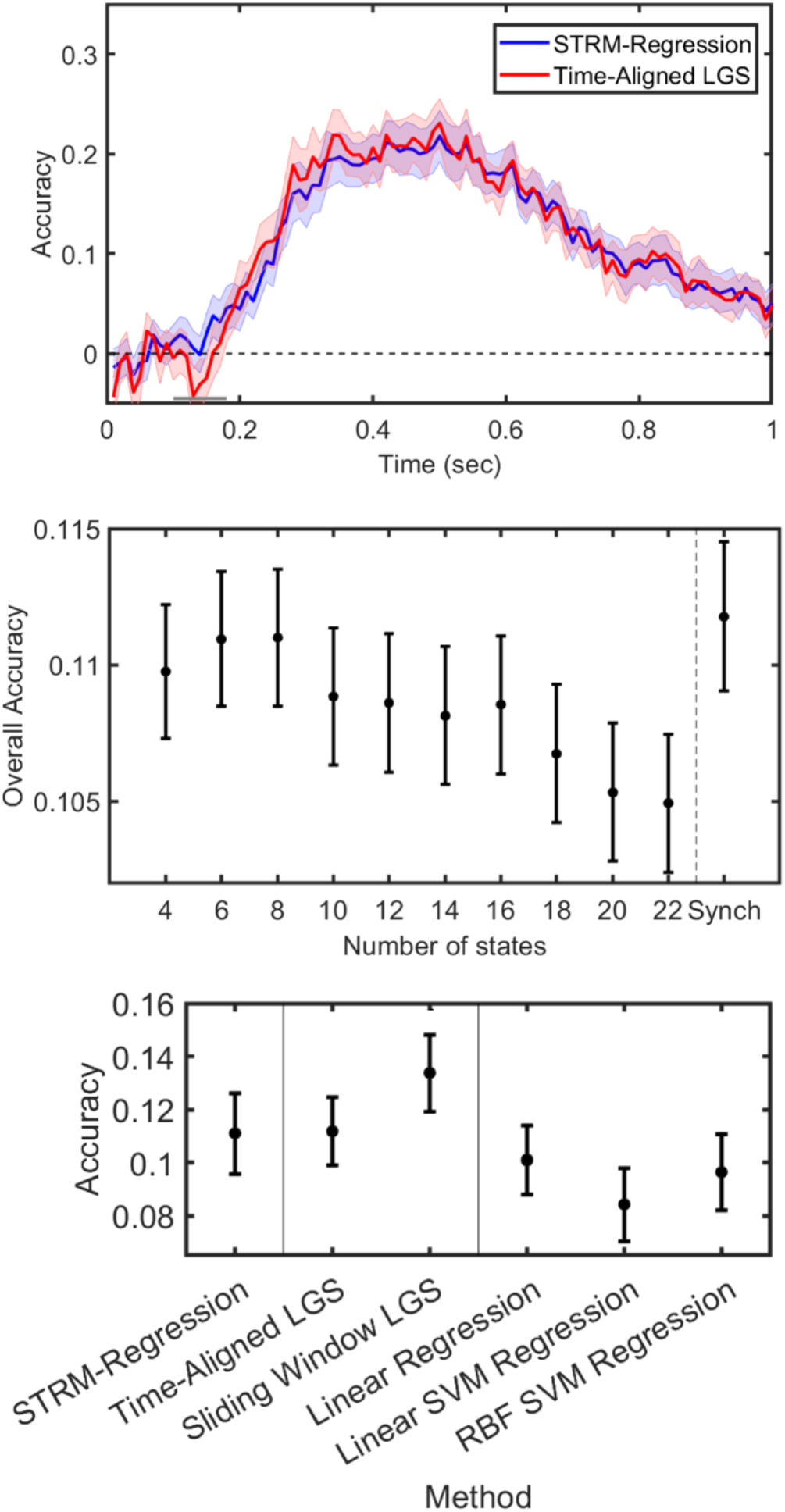
Predictive accuracy of STRM-Regression decoding. Top panel; plotting the Pearson Correlation between model predictions and true regressor values over time, we see equivalent performance by both metrics. The STRM-Regression output shown here was obtained by optimising the value of K (the number of states) by subject-level cross validation (see Methods). Middle panel: this performance is robust over a range of values for the parameter K controlling the number of states, with STRM-Regression displaying no significant difference from synchronous models for all values of K tested (ANOVA, p=0.98); this plot does further identify a performance tuning curve that justifies the use of optimisation through cross validation. Lower panel: Performance of STRM-Regression against a range of different decoding models, both generative (LGS – in both synchronous and optimised sliding window modes) and discriminative (linear regression and Support vector machine regression, using linear and radial basis function kernels). The Pearson correlation shows no significant difference between groups (ANOVA; p=0.19). Whilst not significant, the trend is the same as obtained for STRM-classification: STRM-Regression is broadly consistent in performance with its timepoint-by-timepoint LGS equivalent, however optimised sliding window methods are trending towards superior performance than STRM-Regression. There is no evidence that discriminative models (Linear Regression and SVM regression) in general outperform generative models (LGS and STRM), with the results here trending in the other direction.

We then investigated what drove this performance and how it compared to similar methods. An equivalent encoding model trained using optimised sliding window techniques (ie sliding window LGS; see methods) achieved equivalent performance. There remained a small trend, where the sliding window method performed marginally better than STRM-Regression, mirroring the relationship obtained for STRM-Classification, but these differences were not significant. Overall, they suggest that the model’s sensitivity to trial specific differences in pattern onset timing – whilst showing strong correlations with behavioural variables – is not a major determinant of predictive accuracy. Finally, the question remains whether the overall forward model-based approach itself performs favourably compared to other discriminative methods. To assess this, we compared model performance against that obtained by three decoding methods: linear regression and SVM regression using either linear or radial basis function kernels. Comparing all models on the basis of their correlation coefficient identified no significant group variation (one way ANOVA: p=0.19), despite an apparent trend for the decoding models to achieve lower correlations. Overall, these results suggest that the generative forward model approach to decoding performs no worse than more commonly used regression techniques; and that their use alongside more targeted time series methods (i.e. the STRM and sliding window models) may offer very slight performance improvements.

## 4. Discussion

We have introduced a method for multivariate analysis of task evoked responses that identifies spatially and temporally resolved patterns associated with trial stimuli. We claim that such an approach supports a more direct interpretation of decoding accuracy metrics by linking the classifier predictions to an interpretable linear model of the underlying activity patterns. By resolving these in time, we are able to utilise the high temporal resolution of M/EEG to investigate how the precise timing of activity patterns is modulated over different trials.

Existing methods that assume the same process occurs at the same millisecond on every trial are not just limiting data analysis, but also constraining experimental design. Researchers are currently limited to repetitive paradigms designed to have maximally reproducible timing over trials. We have shown that relaxing this assumption with STRM models reveals meaningful patterns of temporal variation over trials. Ultimately, this new modelling approach could open the door to more flexible experimental designs, allowing tasks more dependent on higher order cognitive functions that have greater variability in their onset timing.

The approach of fitting an interpretable encoding model to training data and then performing a model inversion to make predictions about unseen data is well established in the fMRI literature (Casey et al., 2011; Friston et al., 2008; Kay et al., 2008; Mitchell et al., 2008; Naselaris et al., 2009, 2015; Nishimoto et al., 2011; Schoenmakers et al., 2013), but has only seen limited adoption so far in M/EEG (di Liberto et al., 2015; Kupers et al., 2020). Our focus in this paper has been to emphasise the interpretability benefits of this approach – which are often overlooked – and demonstrate how it can be readily extended to time series analysis for data at high temporal resolution in ways that we believe offer significant benefits to conventional timepoint-by-timepoint decoding.

In particular, we demonstrate that this generative model approach to decoding – with or without the HMM component – achieves equivalent or better accuracies than discriminative classifiers on two different datasets. This corroborates limited evidence from similar surveys of classifiers in M/EEG data (Grootswagers et al., 2017; Guggenmos et al., 2018), suggesting this may be a common feature of this data and the model we propose may generalise well.

Nonetheless, there are a few trade-offs to consider when using the model we have presented. Whilst the observed predictive accuracy was fairly robust across a range of different parameter values, in both datasets any performance gains could be recovered or surpassed by using optimised sliding window techniques, e.g. sliding window LGS in Figure 9 (see SI). There are several possible reasons for this. On the one hand, when computing cross-validated accuracy our methods do not allow an unbiased way to obtain corresponding state timecourses for the held-out test set, instead relying on an estimation procedure that may introduce additional error (see methods). Nonetheless, we did attempt a number of different approaches to estimating state timecourses for held-out data in an unbiased way, none of which consistently outperformed the optimised sliding window techniques (see SI), suggesting that the subtle changes in activity pattern timing are not a major determinant of accuracy. Another potential issue with our model is the assumption of mutual exclusivity between states, which produces sharp jumps in time between inferred activity patterns. If the underlying activity patterns are in fact smooth, then their arbitrary discretization introduces larger errors for timepoints adjacent to state boundaries. On the one hand, this approach is highly interpretable and supports relatively straightforward analyses of the impacts of different behavioural variables on state transition times; on the other hand, this may come at a cost to predictive accuracy around state boundaries. We furthermore expect, given the challenge of overfitting with these datasets that motivated our sequential state constraint, that any comparable dynamic models with a *continuous* state space (eg (Penny & Roberts, 1999)) would be very difficult to suitably regularise.

Our work is very closely related to a number of others; notably a seminal work which showed how any decoding model could be inverted to recover an interpretable forward model of the original data (Haufe et al., 2014). We instead propose the opposite approach; to fit an encoding model of the data and the invert that to make predictions. How are these two approaches different in practice? The first point we would make pertains to the classification accuracies we have reported; when using generative classifiers these are generally either better or at least not significantly different than a range of discriminative models tested, a finding that appears consistent with other examples in the literature (Grootswagers et al., 2017; Guggenmos et al., 2018). Note that all of these discriminative classifiers should, given enough datapoints, converge to the same boundary; generative classifiers however can learn faster if their modelling assumptions are accurate (Ng & Jordan, 2002). This ultimately equates to greater estimation error in the inverted model parameters; that is, the error in forward model parameters obtained via a decoding model inversion is likely to be greater than the error in a directly fitted forward model. Furthermore, the interpretation of an inverted decoding model can be highly problematic wherever regularisation is used (Haufe et al., 2014). Given the ubiquity of regularisation in decoding applications this is not a niche problem. Thus, where interpretation of forward model parameters is the goal, we would argue that one should just fit a forward model directly.

Finally, our work was originally motivated by that of (Vidaurre et al., 2019), which introduced the HMM architecture in the context of decoding models. This demonstrated the potential of time series models for multivariate analysis, but left open the question of model interpretability. Similar to the question we posed in Figure 1, if each state is a different spatial filter, then how should one interpret observed differences in timing on individual trials? Given the lack of direct interpretability of spatial filters themselves, this question does not present an obvious answer. In contrast, by setting up our work around a forward encoding model based on the widely used GLM framework, each state has a clear correspondence to a set of stimulus activation patterns. This affords a straightforward interpretation of each state as a successive stage of stimulus processing, allowing us to then explore its spatial and temporal properties and provide a richer picture of the overall patterns of activation.

## 5. Conclusion

Neuroscientists want to understand what information is represented in brain activity, as well as how and when it is expressed. Existing methods limit what researchers can investigate in several ways. By obscuring the spatial patterns from which decoding accuracy metrics are derived, researchers are often left to interpret a result without clear knowledge of its spatial origins. By assuming the same process occurs at the same millisecond on every trial, researchers are unable to investigate meaningful patterns of temporal variation over trials and are limited in the experimental designs they can pursue.

The STRM model addresses these points with two main innovations; firstly, through the use of an interpretable encoding model to reveal how activity patterns are spatially distributed, and secondly through the use of an HMM to reveal how activity patterns are temporally distributed. M/EEG recordings offer the potential to reveal how the brain represents information at millisecond resolution; the methods we have developed seek to leverage this resolution to simultaneously answer the question of *when* and *where* these patterns emerge. By characterising both the spatial and temporal characteristics of neural representations, we may obtain a more holistic understanding of brain function, offer a new perspective on the role of timing in cognitive processes, and support more flexible experimental designs in the future.

## Author Contributions

C.H.: Formal analysis, investigation, methodology, software, validation, visualization, writing – original draft preparation; D.V.: Conceptualization, methodology, software, supervision, writing – review and editing; N.K.: data curation, investigation, writing – review and editing; Y.L.: data curation, writing – review and editing; T.B.: conceptualization, funding acquisition, supervision, writing – review and editing; M.W.: conceptualization, funding acquisition, methodology, project administration, supervision, writing – review and editing.

## Acknowledgements

The Wellcome Centre for Integrative Neuroimaging is supported by core funding from the Wellcome Trust (203139/Z/16/Z). MW’s research is supported by the NIHR Oxford Health Biomedical Research Centre, the Wellcome Trust (106183/Z/14/Z, 215573/Z/19/Z), the New Therapeutics in Alzheimer’s Diseases (NTAD) study supported by UK MRC and the Dementia Platform UK (RG94383/RG89702) and the EU-project euSNN (MSCA-ITN H2020-860563). TB was supported by Wellcome Trust senior (104765/Z/14/Z) and principal (219525/Z/19/Z) research fellowships, a Wellcome collaborator award (214314/Z/18/Z) and by the James S. McDonnell Foundation (JSMF220020372) during the course of the research. YL was supported by the Max Planck Society. NK’s research is funded by the BBSRC fellowship, BBSRCAFL Fellowship (BB/R010803/1). DV is supported by a Novo Nordisk Emerging Investigator Award (NNF19OC-0054895) and by the European Research Council (ERC-StG-2019-850404). CH is supported by the Wellcome Trust (215573/Z/19/Z).

## Declaration

The authors have no interests to declare.

## Data and code availability

All software code needed to fit the model and run the analyses in this paper is freely available online at https://github.com/OHBA-analysis/HMM-MAR. The two datasets presented in this paper (MEG Visual Stimuli and EEG value based decision making) will be uploaded (upon acceptance) to Mendeley Data and made freely available for download.

## Appendices

### Appendix A: STRM-Classification Bayesian Model inversion

We wish to infer the posterior distribution given by:

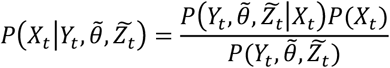

Where 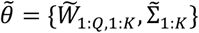 are the maximum a posteriori point estimates learned during training, and 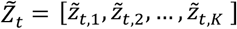 are the state probabilities estimated as outlined in the text. We now introduce the lower case notation *x*_*t*_ = *i* to denote class *i* being active, such that the [1 × *Q*] vector *X*_*t*_ = [*I*(*x*_*t*_ = 1), *I*(*x*_*t*_ = 2), … *I*(*x*_*t*_ = *K*)] where *I* is the Iverson bracket returning 1 if the statement is true and 0 if false. We then have:

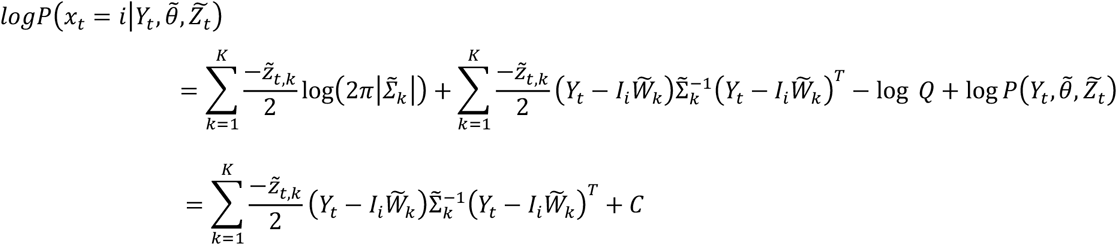

Where the term *C* is constant with respect to the class *i*, and *I*_`_ denotes the [1 × *Q*] vector obtained by taking the *i*th row of the identity matrix of dimension *Q*. As the classes are mutually exclusive and exhaustive, we have

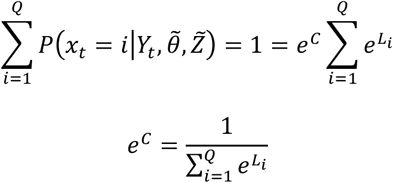

Where *L* is a [*Q* × 1] vector of unnormalized class likelihoods with each entry 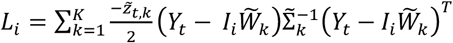. It therefore follows that

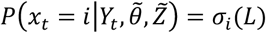

Or in vector notation:

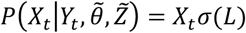

Where *σ*(*L*) is the softmax function which outputs a [*Q* × 1] vector with each entry 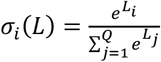.

### Appendix B: STRM-Regression Bayesian Model Inversion

We wish to infer the posterior distribution given by:

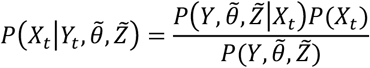

Where *P*(*X*_*t*_)∼*N*(0, *I*_*Q*_) and 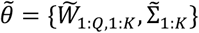 are the maximum a posteriori point estimates learned during training, and 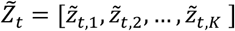 are the state probabilities estimated as outlined in the text. We then have:

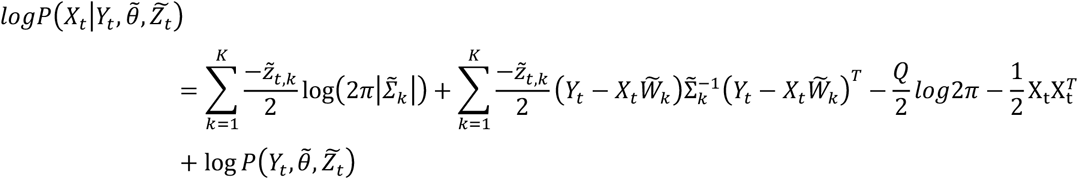

If we make the below substitution:

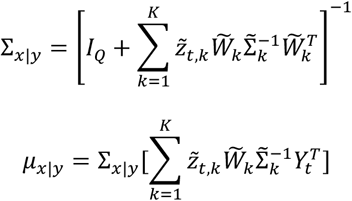

Then the posterior can equivalently be written:

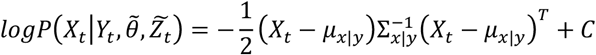

Where the term *C* is constant with respect to *X*_*t*_. This form can be recognised as a normal distribution, equivalently written:

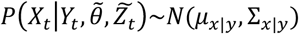

