## Supplementary Info for "Spatiotemporally Resolved Multivariate Pattern Analysis for M/EEG"

**Supplementary information:**

#### 1. Model verification using simulated data

To verify that the model inference procedure functioned as intended, we simulated data from the generative model such that real 'ground truth' parameter values were known; inferred the posterior distribution for these parameters; and verified the accuracy. These are shown below; for further details and other simulations see (Higgins, 2019).

Case 1: STRM-Classification

First we simulated data in two dimensions under two categorical stimuli (along with an intercept term) switching between two sequential latent states ($P = Q = K = 2$). We simulated trials of length T=50 timesteps, with all trials beginning in state 1 and then transitioning to state 2 at some randomly generated timepoint within the trial; transition times were sampled from a uniform distribution on the interval $[2,49]$. We simulated a total of 10 trials, with 5 of each stimulus. Stimulus activation patterns $W_k$ were randomly generated from a standard normal distribution; state covariance patterns $\Sigma_k$ were generated from a Wishart distribution with identity matrix mean and 2 degrees of freedom. Figure S1A plots the data, with red and green colors denoting the class active for that sample. On the right hand side is the ground truth latent state activation simulated for each trial.

Figure S1B shows the inferred state timecourses alongside their true values; these closely match, demonstrating that the model has recovered almost exactly the ground truth state timecourses used to generate the data. Figure S1C plots the data assigned to each latent state, highlighting that these are more clearly separable than the data in figure S1A. Figure S1D shows the mode of the posterior distributions inferred for the model, with cross markers denoting the posterior mean of $W$ and contour plots showing the posterior mode of the covariance matrix $\Sigma$. When compared to the corresponding ground truth model values alongside each plot, they can be seen to match the data and the generative parameter values closely.

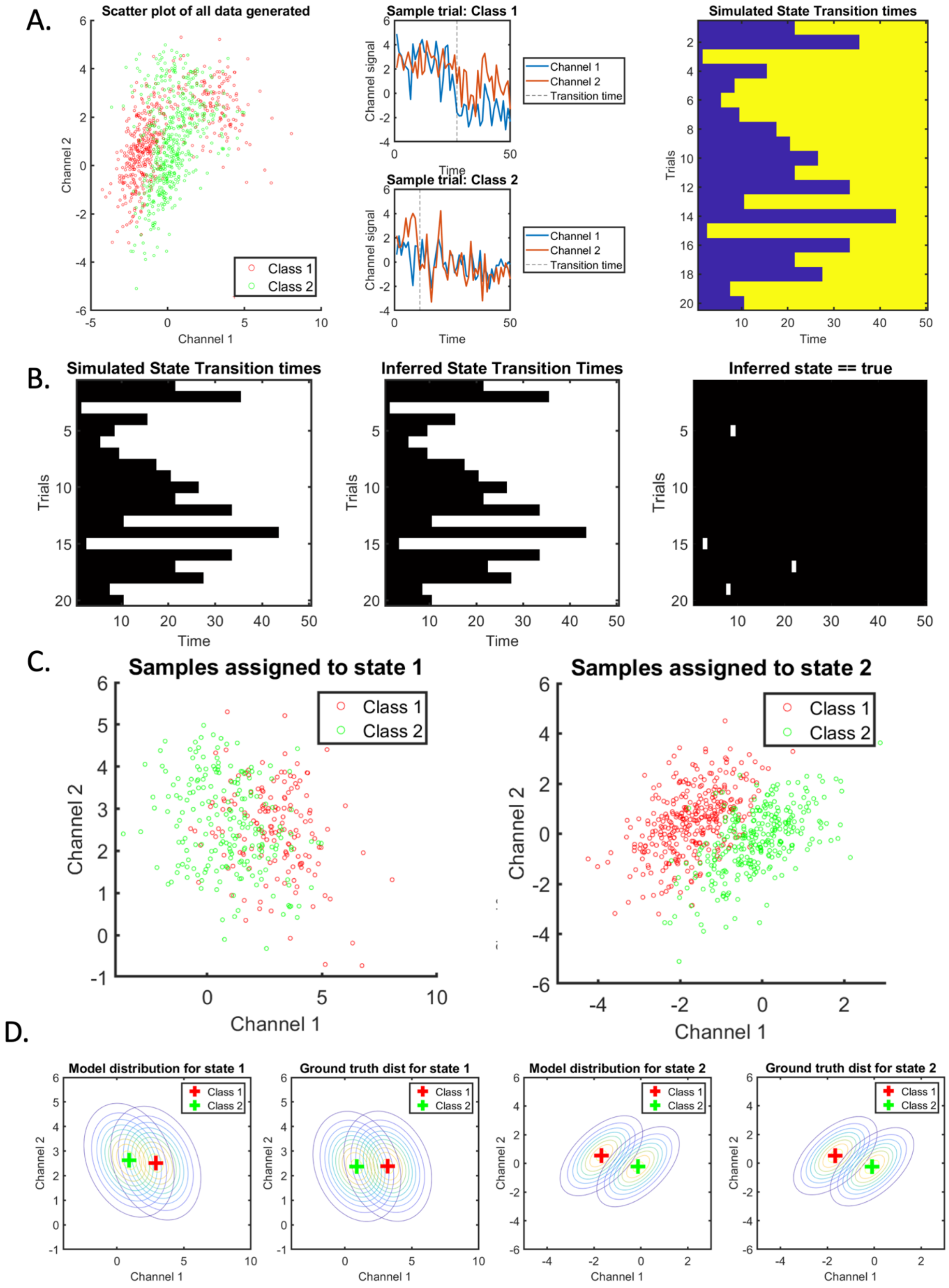


***Figure S1: Ground truth simulations and inferred STRM-Classification model parameters.*** *A. The generated data. We randomly sampled parameters from the generative model as outlined in the text; the scatter plot*
32 *shows the distribution of all datapoints (collapsing over all trials and timepoints), with green and red dots denoting the class the data pertains to. To highlight the temporal evolution, channel data from two sample trials are also plotted against time (one from each class), with the transition time between the two latent*

*states highlighted. B. Inferred state timecourses match simulated ground truth: Plot on the left shows the simulated state timecourses as a raster plot (each row is a trial, each column is a timepoint, and the colouring of white/black denotes that state 1 or 2 respectively is active); the middle plot shows the inferred latent state timecourses, which qualitatively match the ground truth; right plot confirms the inferred latent state matches the ground truth for all but 4 samples (in which the inferred state switch time was out by one timepoint). C. The model separates datapoints into latent states that maximise model fit. Plots show the datapoints assigned into each of latent states 1 and 2 respectively; compared to the scatter plot in A, these are more clearly separable by class. D. Inferred model parameters match ground truth values; plots show the model distribution parameters – i.e. the inferred class mean and covariance – alongside the ground truth parameter values, demonstrating a tight fit.*

Case 2: STRM-Regression

We then simulated from the STRM-Regression model. We again simulated data in two dimensions, with a single continuous valued regressor (along with an intercept term). Regressor values generated for each trial were drawn from a standard normal distribution (the regressor value was fixed for the duration of the trial). We again simulated trials of length 50 samples, with all trials beginning in state 1 and then transitioning to state 2 at some randomly generated timepoint within the trial; transition times were sampled from a uniform distribution on the interval $[2,49]$. We simulated a total of 20 trials. Stimulus activation patterns $W_k$ were randomly generated from a standard normal distribution; state covariance patterns $\Sigma_k$ were generated from a Wishart distribution with identity matrix mean and 2 degrees of freedom. Figure S2A plots the data, with coloured shading indicating the value of the regressor as indicated on the colour bar.

Figure S2B shows the inferred state timecourses alongside their true values; these closely match, demonstrating that the model has recovered almost exactly the ground truth state timecourses used to generate the data. Figure S2C plots the data assigned to each latent state, in each of which a clear and distinct regressor encoding direction and noise pattern is visible. Figure S2D shows the mode of the posterior distributions inferred for the model, with vector arrows denoting the direction and scaling of the regressor encoding direction $W$ and contour plots showing the posterior mode of the covariance matrix $\Sigma$. When compared to the corresponding ground truth model values alongside each plot, they can be seen to match the data and the generative parameter values closely.

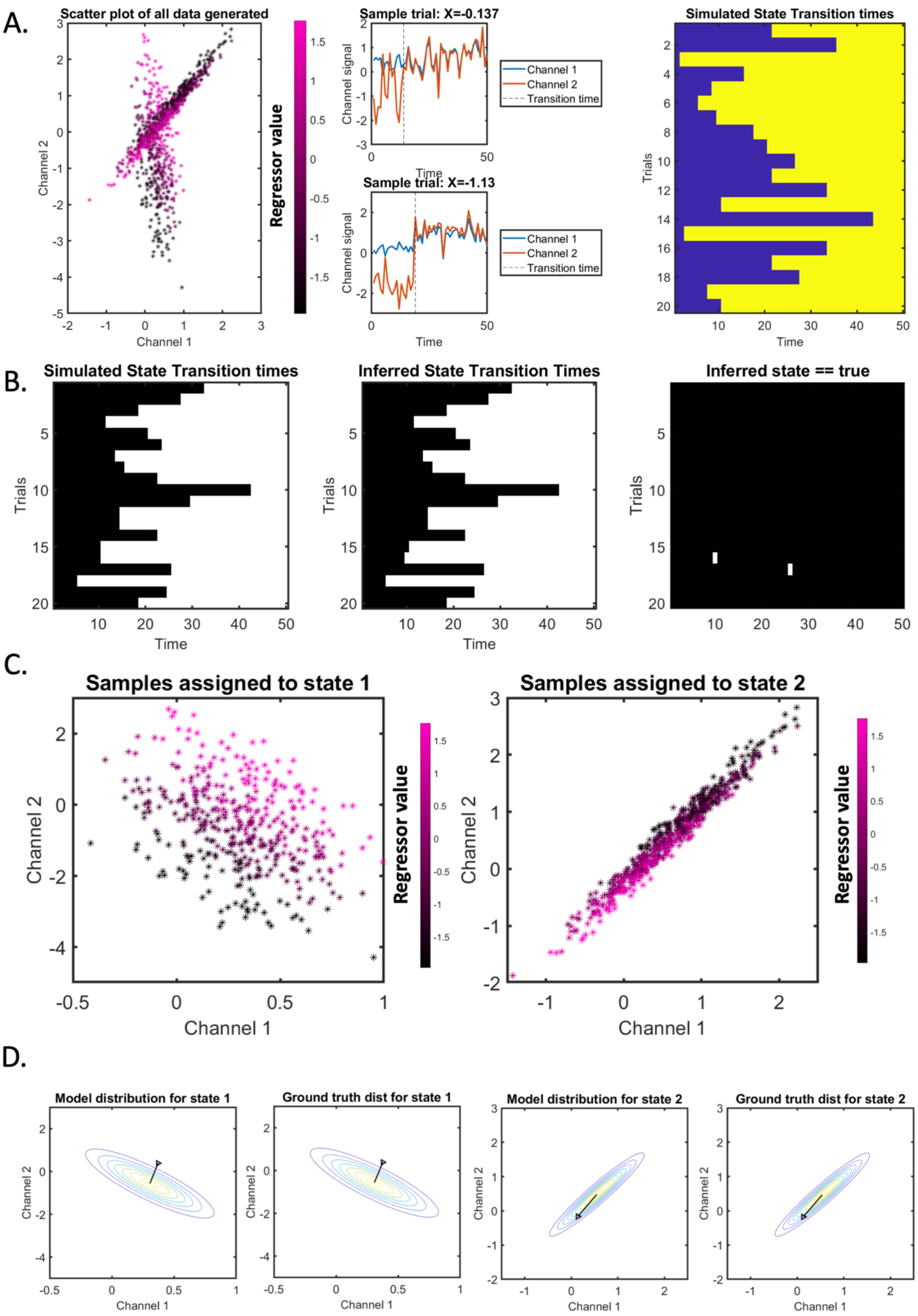


***Figure S2: Ground truth simulations and inferred STRM-Regression model parameters. A.*** *The generated data. We randomly sampled parameters from the generative model as outlined in the text; the scatter plot shows the distribution of all datapoints (collapsing over all trials and timepoints), with colour denoting the value of the sole regressor* $X$ *associated with each datapoint. To highlight the temporal evolution, channel data*

*from two sample trials are also plotted against time (with the regressor value indicated for each), with the transition time between the two latent states highlighted. Right: ground truth simulated state timecourses. B. Inferred state timecourses match simulated ground truth: Plot on the left shows the simulated state timecourses as a raster plot (each row is a trial, each column is a timepoint, and the colouring of white/black denotes that state 1 or 2 respectively is active); the middle plot shows the inferred latent state timecourses, which qualitatively match the ground truth; right plot confirms the inferred latent state matches the ground truth for all but 2 samples (in which the inferred state switch time was out by one timepoint). C. The model separates datapoints into latent states that maximise model fit. Plots show the datapoints assigned into each of latent states 1 and 2 respectively; compared to the scatter plot in A, the distinct linear relationships with the regressor can be more clearly identified. D. Inferred model parameters match ground truth values; plots show the model distribution parameters – i.e. the inferred data covariance and a vector indicating the direction of regressor encoding $W$ – alongside the ground truth parameter values, demonstrating a tight fit.*

#### 2. Test-set state time course fitting

When using cross validation to gauge the model's performance on held-out data (i.e. for the sections computing model predictive accuracy metrics plotted in Figure 6 and Figure 9 of the main text), we face the problem that we cannot know a-priori the correct values of the state timecourses for the held out test set; the STRM model differs from standard MVPA approaches in that these model parameters are trial-specific.

Whilst we are unable to infer these parameters directly using the model, we can resort to procedures that estimate them in a principled and unbiased way. As outlined in the text we used a linear regression model to estimate the state timecourses from the data directly. We explain this method in more detail here and justify this choice over several alternatives.

Specifically, as in Figure S3, the cross validation procedure involves (i) partitioning the data into training and test folds; (ii) learning all STRM model parameters from the training set; (iii) computing the expected value of the observation model (ie non-trial-varying) parameters $\widetilde{W}_{1:K}$ and $\tilde{\Sigma}_{1:K}$ from the posterior; (iv) estimating a linear relationship between the training data latent state parameters (ie the parameters that are unique to each trial: $\tilde{z}_t$ for $t$ in the training set); note we train a different model for each timepoint within the trial (v) using this linear model to estimate the test set latent state parameters (ie $\tilde{z}_t$ for $t$ in the test set); (vi) computing the predictions of the model for the test set and computing the accuracy of these, and (vii) repeating for each cross validation fold.

We did investigate several different methods for estimating the latent state parameters $\tilde{z}_t$ from the data, which we briefly justify here: overall, we found that the regression model was robust over different data types, and fast (an important consideration given the number of folds over which these computations had to be repeated). In figure S3, we compare the following methods:

(i) The regression method outlined in the main text and above;

(ii) Computing the *mean state timecourse* $\tilde{z}_t$ observed over trials in the training set as the each test set trial's state timecourse. Note that in this case there are no inter-trial differences in the encoding model fit as the mean over trials does not vary.

(iii) Learning an *equivalent unsupervised model*. This procedure involves holding the inferred state timecourses for the training data fixed and learning the parameters of an unsupervised model of the type used in (Vidaurre et al., 2016); specifically, inferring a single mean and covariance across all channels of the data for each state, with no knowledge of the design matrix. Such a model could then be fit to the test set data to infer state timecourses in an unbiased way.

As an example, Figure S4 shows the accuracy achieved fitting a STRM model with K=12 states, using each of these methods to estimate the held-out test set state timecourse $\tilde{z}_t$. Across the three different estimation methods evaluated, using the mean state timecourse achieves slightly better results than using the regression model; whilst using the equivalent unsupervised model achieves significantly poorer performance.

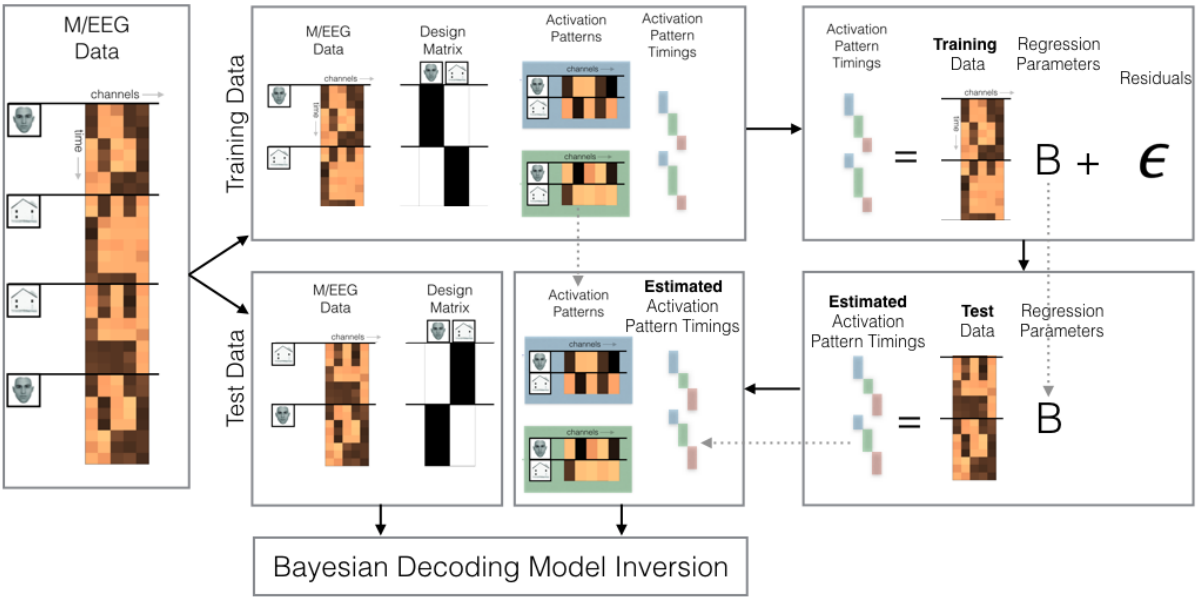


***Figure S3:*** *The full cross validation procedure. Cross validation involves partitioning the data into training folds and test folds (left hand side), and training the STRM model on the training data. The STRM model includes some parameters that can be used directly on the test set (the activation patterns) and some parameters that are unique to each trial (the activation pattern timings) and therefore cannot directly be applied to the unseen trials in the test data. We therefore apply a post-hoc procedure (right hand side of diagram) to estimate a suitable set of activation pattern timings for the test set in an unbiased way. This applies training a linear regression model (top right) to estimate a relationship between the training data and its corresponding activation pattern timings. Note that we train a distinct set of regression weights for each timepoint within the trial. We then apply these regression weights to the test data to obtain estimated activation pattern timings*

*(bottom right). These estimated activation pattern timings can then be used in combination with the previously learned activation patterns to make predictions via the Bayesian decoding model inversion outlined in the text.*

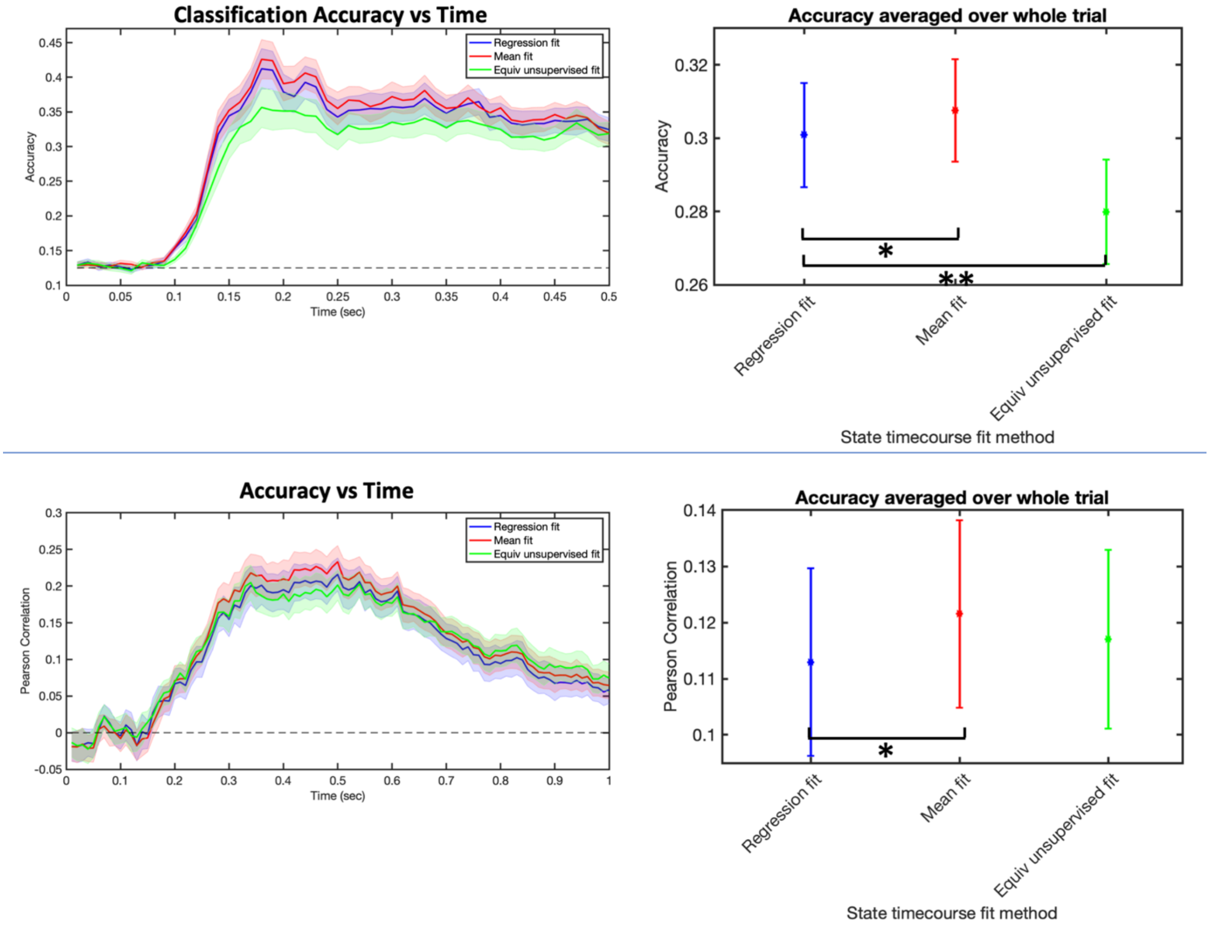


***Figure S4**: Comparing accuracy of different cross validated state fitting methods. Top panel: applying different methods to the STRM-Classification paradigm and measuring the classification accuracy (mean over subjects +/- ste), either versus time (left) or averaged over all timepoints (right). Using the mean fit method provided a slight improvement on the regression fit method; using the equivalent unsupervised fit method was significantly worse. A similar result was obtained on the STRM-regression paradigm: we find an improvement when using the mean fit method over the regression fit method; and no significant difference between the regression and equivalent unsupervised fit methods.*

Fitting an equivalent unsupervised model tests whether the same information that determines stimulus activation pattern timings in the STRM model could be reflected more generically with a dynamic model that is common over all stimuli. The relatively poorer performance of the unsupervised HMM model confirms the STRM model is indeed using stimulus specific information in determining state activation timings, and that this timing information is likely reflected in distributed subtle patterns of variation that are not well reflected by unsupervised modelling.

Applying the mean state timecourse effectively ignores any time-varying information in the held out test set, applying a common pattern across all trials. The slightly improved performance achieved by doing so – especially when taken together with the improved performance of optimised sliding window methods presented in section 3.1.3 and 3.2.3 – suggests either that the methods we have used in cross validation are outputting a poor estimate of state timing information, or that this information is not actually that helpful for classification accuracy. We should note that in our own experimentation we tried quite an exhaustive list of alternative methods for state timing estimation, none of which consistently outperformed the mean fit on the above metrics. Furthermore, as outlined in the main text, the gain in accuracy shown by the HMM model can equivalently be achieved and for some cases overcome by optimised sliding window techniques, that by design have no sensitivity to timing difference across different trials. Thus, as outlined in the text, we have tentatively concluded that knowledge of exact state timing information – which we have shown to reliably correlate with behavioural variables – is not actually particularly informative for improving classification accuracy. There are a number of reasons why this could be the case.

Firstly, the mutual exclusivity assumption imposed by the STRM model is a very strong assumption. It is methodologically quite useful, allowing characterisation of successive stages of processing, but discretising brain activity in this way is potentially counterproductive when assessing classification accuracy over an entire trial. Importantly, when fitting the mean state timecourse, one fits a smoothly averaged mixture of different states at any point in time – transitions between states tend to be much smoother than in the alternative regression fit model – which could achieve more consistent performance if the underlying neural activity trajectories are themselves smoothly varying rather than discrete. Secondly, it is possible simply that the signal to noise ratios are not sufficient to estimate these state timecourses on held out trials. The poor performance of the equivalent unsupervised models suggest the information for these timings is distributed subtle patterns of variation – it is possible that these are simply below some threshold needed to be sufficiently estimated from the data without knowledge of the stimulus.

Finally, one may ask why we have used the regression fit model in the text if the mean fit in fact achieved a better performance. Given all other analyses focus on the time-varying patterns in the data, we wanted to be very clear on the accuracy associated with such time-varying estimates, which is best reflected by the regression fit model. As discussed above, the mean fit imposes the same state timecourse over all trials, and thus does not reflect the influence of time varying dynamics on predictive accuracy.

### Supplementary Information – References


Higgins, C. (2019). Uncovering temporal structure in neural data with statistical machine learning models. In *Doctoral Thesis*. University of Oxford.

Vidaurre, D., Quinn, A. J., Baker, A. P., Dupret, D., Tejero-Cantero, A., & Woolrich, M. W. (2016). Spectrally resolved fast transient brain states in electrophysiological data. *NeuroImage*, *126*, 81–95. https://doi.org/10.1016/j.neuroimage.2015.11.047
